# Full assembly of HIV-1 particles requires assistance of the membrane curvature factor IRSp53

**DOI:** 10.1101/2021.02.10.430663

**Authors:** Kaushik Inamdar, Feng-Ching Tsai, Aurore de Poret, Rayane Dibsy, John Manzi, Peggy Merida, Remi Muller, Pekka Lappalainen, Philippe Roingeard, Johnson Mak, Patricia Bassereau, Cyril Favard, Delphine Muriaux

## Abstract

During HIV-1 particle formation, the requisite plasma membrane curvature is thought to be solely driven by the retroviral Gag protein. Here, we reveal that the cellular I-BAR protein IRSp53 is required for the progression of HIV-1 membrane curvature to complete particle assembly. Partial gene editing of IRSp53 induces a decrease in viral particle production and a viral bud arrest at half completion. Single molecule localization microscopy at the cell plasma membrane shows a preferential localization of IRSp53 around HIV-1 Gag assembly sites. In addition, we observe the presence of IRSp53 in purified HIV-1 particles. Finally, HIV-1 Gag protein localizes preferentially to IRSp53 I-BAR domain induced curved membranes on giant unilamellar vesicles. Overall, our data reveal a strong interplay between IRSp53 I-BAR and Gag at membranes during virus assembly. This highlights IRSp53 as a crucial host factor in HIV-1 membrane curvature and its requirement for full HIV-1 particle assembly.

## Introduction

The cell plasma membrane is a dynamic organelle, where crucial processes such as endocytosis and exocytosis take place through local membrane deformations. Several pathogens, such as bacteria and enveloped viruses interplay with the plasma membrane in the course of their replication cycle. Pathogens often enter the cells by endocytosis (1, 2) and exit by membrane vesiculation (3, 4), which are processes linked to generation of plasma membrane curvature; either inward or outward deformations. HIV-1 is an enveloped positive strand RNA virus belonging to the family Retroviridae, known to assemble and bud outward from the host cell plasma membrane (5). The structural Gag polyprotein of HIV-1, by itself, is responsible for particle assembly (6): It can oligomerize at the inner leaflet of the plasma membrane forming virus-like particles (VLPs). The force required to bend the membrane to achieve VLP formation has been proposed to be provided by Gag self-assembly (7). The self-assembly of Gag has also been recently shown to segregate specific lipids (8, 9) and proteins (10), generating plasma membrane domains that could favor budding (11, 12). However, only a small proportion of initiated clusters of Gag reaches the full assembly state leading to VLP release in living CD4 T cells (13), a majority being aborted events. Therefore, the mechanism by which the virus overcomes the energy barrier associated with the formation of the full viral bud remains an open question. Recently, coarse grained model of HIV assembly has shown that self-assembly of Gag might not be sufficient to overcome this energy barrier (14) leaving the assembly in intermediate states supporting the fact that other factors may be necessary to assist Gag self-assembly during the generation of new VLPs. Indeed, the plasma membrane curvature can be also generated by diverse host cell proteins. For example, I-BAR domain proteins sense and induce negative membrane curvature at a few tens to one hundred of nanometer-scale, i.e. in the HIV-1 particle diameter size range, while generating outward micrometer-scale membrane protrusions such as membrane ruffles, lamellipodia, and filopodia. IRSp53 was first discovered as a substrate phosphorylated downstream of the insulin receptor (15). It is also the founding member of membrane curving I-BAR domain protein family, whose other mammalian members are MIM (missing-in-metastasis), ABBA (actin-bundling protein with BAIAP2 homology), PinkBAR (planar intestinal and kidney specific BAR domain protein), and IRTKS (insulin receptor tyrosine kinase substrate)(16). In addition to interactions with plasma membrane, IRSp53 binds both Rac1 through its N-terminal I-BAR domain (17) and Cdc42 directly through its the unconventional CRIB domain (18), as well as downstream effectors of these GTPases such as WAVE2, Mena, Eps8 or mDia, through the SH3 domain. Thus, IRSp53 functions as a scaffold protein for the Rac1/Cdc42 cascade (19). IRSp53 was reported to exhibit a closed inactive conformation that opens synergistically upon binding to Rac1/Cdc42 and effector proteins (20–23). Regulation of IRSp53 activity was recently shown to occur through its phosphorylation and interaction with 14-3-3 (24). Structurally, the I-BAR domain of IRSp53 is composed of crescent-shaped rigid six alpha-helix bundle dimer. Due to its concave membrane binding surface and lipid interactions, IRSp53 is able to generate negative membrane curvature (16). While capable of forming homo-dimers, IRSp53 is also able to recruit and form hetero-dimers with other proteins to form clusters for initiation of membrane curvature (20). Since the Rac1/IRSp53/Wave2/Arp2/3 signaling pathway is involved in the release of HIV-1 particles (25), we hypothesized that IRSp53 may be a prime candidate for membrane remodeling required during viral bud formation. Hence, we investigated the possible role of IRSp53 and its membrane curvature generating activity in HIV-1 Gag assembly and particle budding. Importantly, we discovered that IRSp53 is present in an intracellular complex with HIV-1 Gag at the cell membrane, incorporated in Gag-VLPs and associated with purified HIV-1 particles, and that IRSp53 functions in HIV-1 assembly as a facilitator of optimal HIV-1 particle formation through its membrane bending activity. Thus, we identify IRSp53 as an essential non-redundant novel factor in HIV-1 replication, and demonstrate that it is critical for efficient HIV-1 membrane curvature and full assembly at the cell plasma membrane.

## Results

### IRSp53 knockdown decreases HIV-1 Gag particle release by arresting its assembly at the cell plasma membrane

We report here that partial knockdown of IRSp53 reduces HIV-1 particle release in host Jurkat T cells and in model cell line HEK293T (Fig1a, b), as we also reported previously in primary T lymphocytes (Thomas et al., 2015). Cells were treated with siRNA targeting IRSp53 or IRTKS (validated by extinction of the transfected ectopic IRSp53-GFP or IRTKS-GFP proteins – Fig S1b and S1c respectively). In the context of Jurkat T cells, when expressing the viral Gag proteins in the context of HIV-1(ΔEnv) in order to only monitor the late steps of the viral life cycle, the partial IRSp53 knockdown (maximum at 50% of gene extinction) reduced particle release by 3 fold as compared to the control siRNA (Fig.1a, Left), and reached 6 fold by calculation if reported to the percentage of gene extinction.

**Fig. 1.**
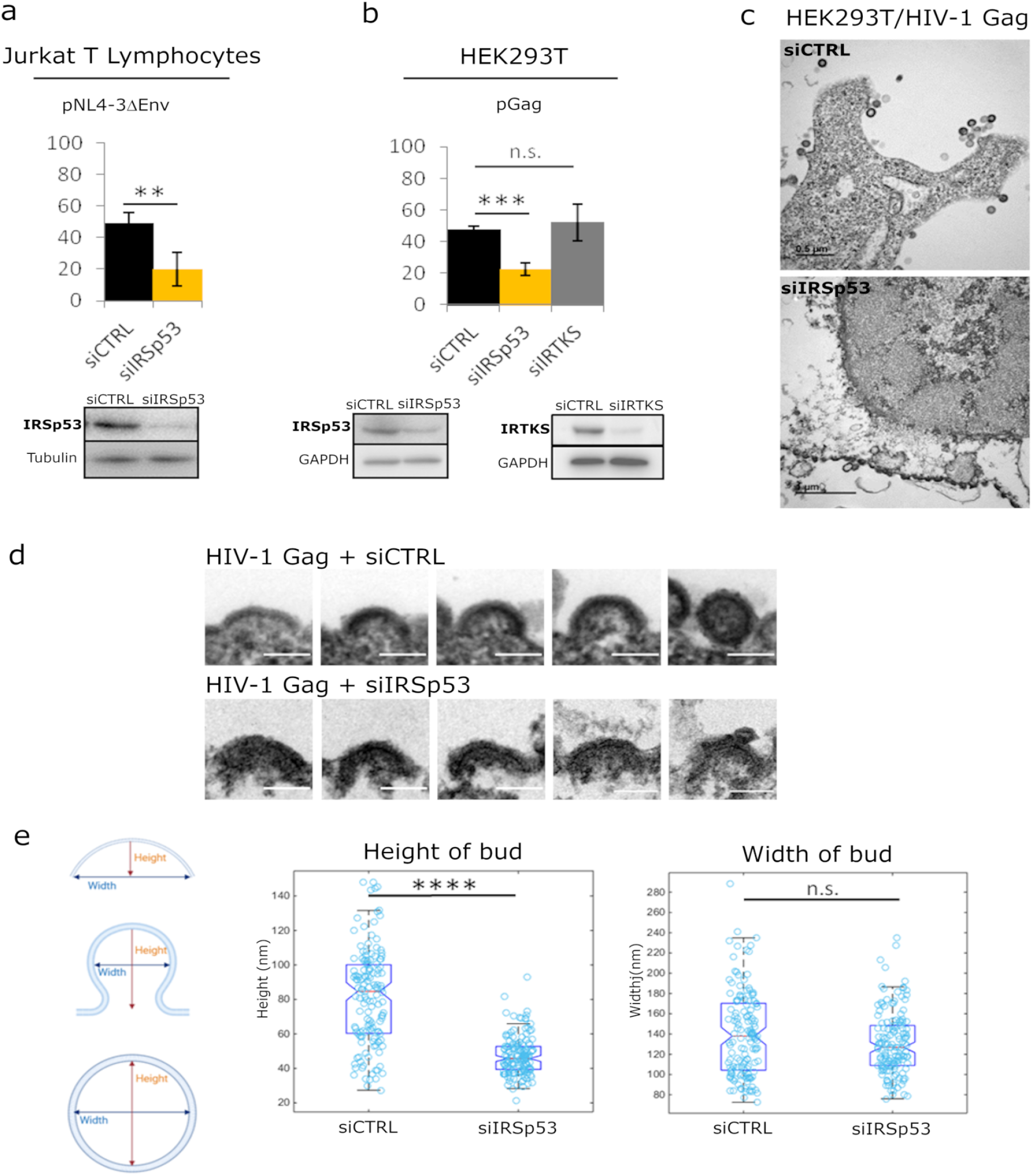
Partial knockdown of IRSp53 decreases HIV-1 Gag particle release by arresting assembly at the cell plasma membrane. a) siRNA knockdown of IRSp53 in Jurkat T lymphocytes leads to a significant decrease in pNL4-3**Δ**Env particle release (left graphs, immunoblots for IRSp53, p=0.00265, Student’s t-test, and loading controls beneath the graphs). Similarly, knockdown of IRSp53 in HEK293T cells led to a significant decrease in HIV-1 Gag particle release (p=0.00487, Student’s t-test). On the opposite, knockdown of a closely related I-BAR protein IRTKS did not have a significant effect on particle release (p=0.092463, Student’s t-test, right graphs, immunoblots for IRSp53, IRTKS and loading controls beneath the graphs) (n=3 independent experiments). c) Transmission electron microscopy images of HEK293T cells expressing HIV-1 Gag with control siRNA (upper panel) and siRNA IRSp53 (lower panel). d) Cells knocked down for IRSp53 show arrested buds at the plasma membrane as compared to the control cells which display a normal range of buds in different stages of assembly and budding (scale bar is 100 nm). e) Measurement of the bud dimensions (height and width median with interquartile) in the control siRNA and siRNA IRSp53 conditions (n=145 buds from 14 different cells for each condition). The knocked down cells exhibit a narrow range of heights corresponding to the arrested buds visible in the images while the control cells display a wider range of heights corresponding to assembly progression (left graph, “Height of bud”). Distribution of the height values in the two conditions are significantly different (p=1.05×10^−28^, Kolmogorov-Smirnov test). On the opposite, the widths of the buds in both conditions did not display significant differences in distributions. (p=0.0609, Kolmogorov-Smirnov test).

This reduction in HIV-1 particle release is highly significant (n=3 independent experiments, p value = 0.00265, Student’s t-test) since the gene editing of IRSp53 cannot be complete, nor edited by CRISPR/Cas9 knockout, without being toxic for the cells, which renders this study particularly delicate timewise. To compare the role of different I-BAR domain proteins from the same family, we also measured the effect of siRNA targeting IRSp53 and IRTKS (Fig.1b) on HIV-1 Gag virus-like particle (VLP) production in HEK293T cells (see graph Fig.1b, and immunoblots Fig. S1d, e). IRTKS shares similar protein domain organization and high sequence homology with IRSp53 (40% amino acid sequence identity and 59% sequence similarity, Fig. S2), and displays some functional redundancy with IRSp53 (26). IRTKS can also induce plasma membrane curvature (27). Partial knockdown of IRSp53 (50% gene inhibition) resulted in a 2-3 fold decrease in HIV-1 Gag particle production (n=3 independent experiments, p value = 0.000487, Student’s t-test) (Fig.1b, Right, Fig. S1d). In contrast, knockdown of IRTKS (Fig.1b, Fig. S1e) did not have any significant effect on HIV Gag particle release (n=3 independent experiments, p value = 0.092463, Student’s t-test), thus precluding the possibility of redundant functions between IRSp53 and IRTKS in the context of HIV-1 Gag particle formation. Electron microscopy imaging of siRNA IRSp53 treated HEK293T cells expressing HIV-1 Gag revealed particle budding arrests at the cell plasma membrane (Fig.1c, Lower Panel), as compared to the siRNA-control cells (Fig.1c, Upper Panel). While the control cells exhibited the normal phenotype of Gag-VLP budding from the cell plasma membrane, the IRSp53 knockdown cells displayed a series of viral buds arrested in assembly decorating the cell plasma membrane (Fig.1c, Fig S3). These results revealed an arrest in Gag assembly at the membrane and thus the involvement of IRSp53 in the assembly process. Since IRSp53 is an I-BAR protein involved in cell membrane curvature, we measured the curvature exhibited by HIV-1 buds in IRSp53 knockdown cells. While control cells displayed a range of HIV-1 Gag particles at different stages of assembly and budding, cells knocked down for IRSp53 displayed arrested buds at an early assembly stage (Fig.1d). On measuring the dimensions of these arrested buds, we found that buds from cells knocked down for IRSp53 displayed a narrower range of curvature height (48 ± 22nm) as compared to the control (85 ± 53 nm)(n=145 buds from 14 different cells, p value = 1.0539 x 10^-28^ Kolmogorov-Smirnov Test), while the bud widths presented no difference between siIRSp53 (135 ± 64nm) and the control (140 ± 87 nm) (n=145 buds from 14 different cells, p value = 0.0609, Kolmogorov-Smirnov Test) (Fig.1e). The control cells thus exhibited a range of height and widths consistent with the range of buds seen at the membrane of these cells. The result indicates that in the absence of IRSp53, the viral buds were unable to progress beyond a certain curvature.

### Complexing and increase in membrane binding of IRSp53 upon HIV-1 Gag expression in cells

Since both Gag and IRSp53 target the cell plasma membrane upon interaction with PI(4,5)P2 (9, 10, 27–31), we then tested if Gag and IRSp53 could associate directly or indirectly using immuno-precipitation assays (Fig.2). Our results showed that immuno-precipitation of endogenous IRSp53 resulted in co-precipitation of Gag (Fig.2a, lane 1) as compared to controls (lanes 2 to 4). Unfortunately, we could not access the amount of IRSp53 pull down by the antibody between conditions because the IgG signals completely masked endogenous IRSp53. To overcome this issue, we performed the same experiment with ectopic IRSp53-GFP confirming the pull down of IRSp53-GFP and Gag by the IRSp53 antibody (Fig. S4a). We concluded that HIV-1 Gag and IRSp53 were components of the same intracellular complex, interacting directly or indirectly through other factors or membrane domains. IRSp53 is a cellular protein that switches from the cytosol to the cell plasma membrane for inducing membrane ruffles upon activation by Rac1 and its effectors (22, 32).

**Fig. 2.**
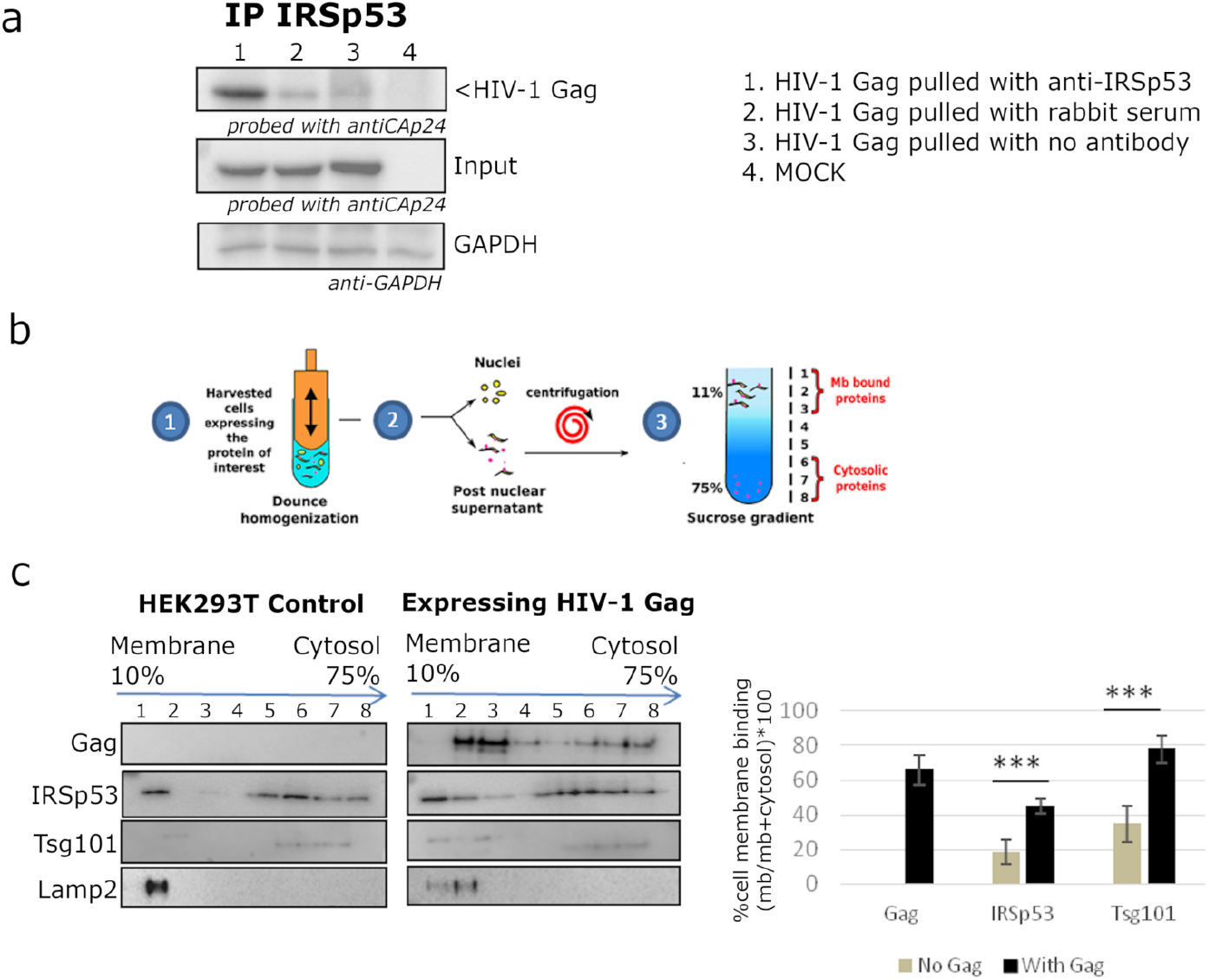
HIV-1 Gag and IRSp53 complexing and cell membrane bindings. HIV-1 Gag and IRSp53 complexing and cell membrane bindings. a) Immunoprecipitation of HIV-1 Gag with an anti-IRSp53 antibody. HIV-1 Gag is enriched in anti-IRSp53 pulldown, as compared to the controls. b) Membrane flotation assay protocol: (1) 293T cells were dounced, (2) the post-nuclear supernatant was loaded on a discontinuous sucrose gradient, and (3) following ultracentrifugation, cell membranes were separated from the cytosolic fraction. c) Immunoblots of the indicated proteins (left panel) and quantification of the % of protein membrane binding in the graph (right panel) show that upon Gag expression in cells, IRSp53 is significantly enriched by 2-fold in the cell membrane fraction (p=0.001753, Student’s t-test) (n=5 independent experiments). A similar increase is observed for Tsg-101, a known interactor of the Gag-p6 polyprotein (p= 0.001974).

We have previously shown that Gag cellular expression triggers Rac1 activation (25), on which IRSp53 membrane localization and function is dependent. Thus, here, we compared the relative membrane binding of IRSp53 upon cellular expression of HIV-1 Gag using membrane flotation assays (Fig.2b). In the absence of Gag (“HEK293T control cells”), one could observe the presence of IRSp53 both in the cytosol (fractions 6-8) and at the cell membranes (fractions 1-2). The lysosomal membrane protein Lamp2 was used to indicate the membrane fraction. Thus, at equilibrium, 19 ± 8% of IRSp53 was bound to cell membranes (Fig.2c). The same experiment was repeated with cells expressing HIV-1 Gag, where 66 ± 9% of Gag was bound to the cell membranes (Graph, Fig.2c). Notably, we observed a 2-fold increase with 44 ± 5% of IRSp53 bound to the cell membrane upon HIV-1 Gag expression (Fig.2c)(n=5 independent experiments, p value = 0.001735, Student’s t-test). Interestingly, we observed the same increased membrane binding (1.5 fold) of IRSp53 I-BAR in the presence of HIV-1 Gag on giant unilamellar vesicles (GUVs) (Fig. S11 a,b). This effect was comparable with the one of Tsg101, a protein of the ESCRT-I complex known to interact mainly with the p6 domain of Gag (33–35). The cellular endosomal sorting complex required for transport (ESCRT) machinery has been involved into the mechanism of vesicular budding of intracellular multi-vesicular bodies, and also hijacked by the HIV-1 Gag protein for viral particle budding. Here, we observe a 2-fold increase in cell membrane binding of Tsg101 upon Gag expression, passing from 36 ± 10% without Gag to 79 ± 8% in the presence of Gag (Fig.2c))(n=3 independent experiments, p value = 0.001974, Student’s t-test). Furthermore, we examined if Gag/IRSp53 complexing was dependent on the p6 domain of Gag to reveal if this could be independent of ESCRT recruitment by Gag. We thus used a C-terminal mutant of Gag, GagΔp6, which is deficient in ESCRT-Tsg101 recruitment (35), still capable of binding the plasma membrane and assembling particles, but that buds poorly. Thus, GagΔp6 viral particles were tethered and stay attached to the plasma membrane (see (13) for the characterization of Gag(i)mEos2Δp6). Our experiments revealed that Gag, Gag(i)mEos2 and GagΔp6(i)mEos2 were all pulled down with IRSp53 (Fig S4), showing that addition of the internal mEos2 protein did not affect the complexing of Gag with IRSp53, which tag is required for super resolution microscopy imaging of Gag (see following section). Moreover, we showed that the p6 domain was not required for Gag/IRSp53 molecular interplay. Together, these results suggest that there is a complexing between HIV-1 Gag and IRSp53 reinforcing the idea of a strong molecular interplay between these two proteins directly or indirectly but in the same membrane domain. We evidenced that cellular Gag expression, most probably by triggering Rac1 activation (25), favors cell membrane binding of IRSp53.

### Single Molecule Localization Microscopy reveals IRSp53 surrounding HIV-1 Gag assembly sites

Our finding that IRSp53 and HIV-1 Gag are present in the same molecular complex at the cell membrane motivated us to assess whether IRSp53 was present specifically at the Gag assembly sites. Because HIV-1 assembly are ~ 100 nm in diameter (13, 36), we used PALM (Photo-Activated Localization Microscopy) coupled to dSTORM (direct Stochastic Optical Reconstruction Microscopy) with TIRF illumination, to investigate with high resolution the localization of I-BAR proteins in Gag(i)mEos2 assembly sites at the plasma membrane. Using density based spatial scan (DBSCAN) of Gag localizations, we quantified the size distribution of the Gag clusters, at a localization precision of ~16 nm (Fig S5), and we found a diameter of 80 to 100 nm, which is within the size-range of HIV-1 Gag assembly sites (13, 36) (Fig.3a,b). Reconstructed dual color PALM/STORM images exhibited Gag(i)mEos2 assembly sites close to or overlapping IRSp53 (Fig.3a, Fig.S6) whereas Gag clusters did not seem to overlap with IRTKS (Fig.3b, Fig.S6), consistent with the results of siRNA presented in Fig.1b. In order to quantify these observations, we performed coordinate-based colocalization (37) (CBC) analysis of HIV-1 Gag and IRSp53 (or IRTKS) (Fig. S7 for the process workflow). In contrast to classical colocalization analysis, CBC takes into account the spatial distribution of biomolecules to avoid excessive colocalization due to local densities and provides a colocalization value for each single-molecule localization. This CBC value ranges from −1 to +1, where −1 corresponds to anti-correlation, 0 indicating non-correlation and +1 corresponds to perfect correlation between the two molecules. Since CBC values are calculated for each localization, we plotted the CBC values as cumulative frequency distributions for all localizations (n>105) of IRSp53/Gag and IRTKS/Gag. As shown in Fig.3c, the CBC distribution for IRSp53/Gag has a higher proportion of values exhibiting high colocalization (27% of CBC>0.5) in comparison with IRTKS/Gag values (14% of CBC>0.5) which on the opposite show a very high proportion (close to 75%) of anti-correlation (CBC<0) (Fig.3d). This comparison directly shows that IRSp53 displays stronger single molecule colocalization with Gag in assembling clusters than IRTKS does. However, the average positions of IRTKS or IRSp53 molecules with respect to Gag molecules within the assembling clusters were unclear. Thus, we performed simulations to generate different patterns of PALM/STORM localizations (Fig. S8). Comparison of the CBC values of simulated data with our experimental values indicated that IRSp53 localization, on average, displays a restricted pattern around and in the assembly sites. This corresponds to a circular ring surrounding the assembly site at 80 nm from the center of the Gag budding sites with a width of 80 nm (Fig.3e, Fig. S8e). On the other hand, IRTKS was present as a large diffuse pattern centered at 140 nm from the Gag assembly site center with a width of 200 nm, explaining why fewer IRTKS molecules were detected in the assembly sites (Fig.3f, Fig. S8f). Our results thus show that IRSp53 indeed specifically localizes at HIV-1 Gag assembly sites at the cell plasma membrane, whereas IRTKS poorly does. Finally, the same imaging (Fig. S9) and CBC analyses (Fig. S10) were applied to the HIV-1 host CD4 T cells, i.e. Jurkat T cells expressing Gag(i)mEos2 and immuno-labelled for IRSp53 (Fig S10a) or for IRTKS (Fig S10b). The results showed that the CBC cumulative distributions for IRSp53 (Fig S10c,e), as well as for IRTKS (Fig S10d,f), are similar to those obtained in HEK293T cells (Fig.3c,d). These analyses confirm the preferred IRSp53 localization to Gag assembly site in the host CD4 T cells. The involvement of IRSp53 around Gag assembly sites seems to be conserved regardless of the cell type, reinforcing the idea of a specific role for IRSp53 in HIV-1 Gag particle assembly.

**Fig. 3.**
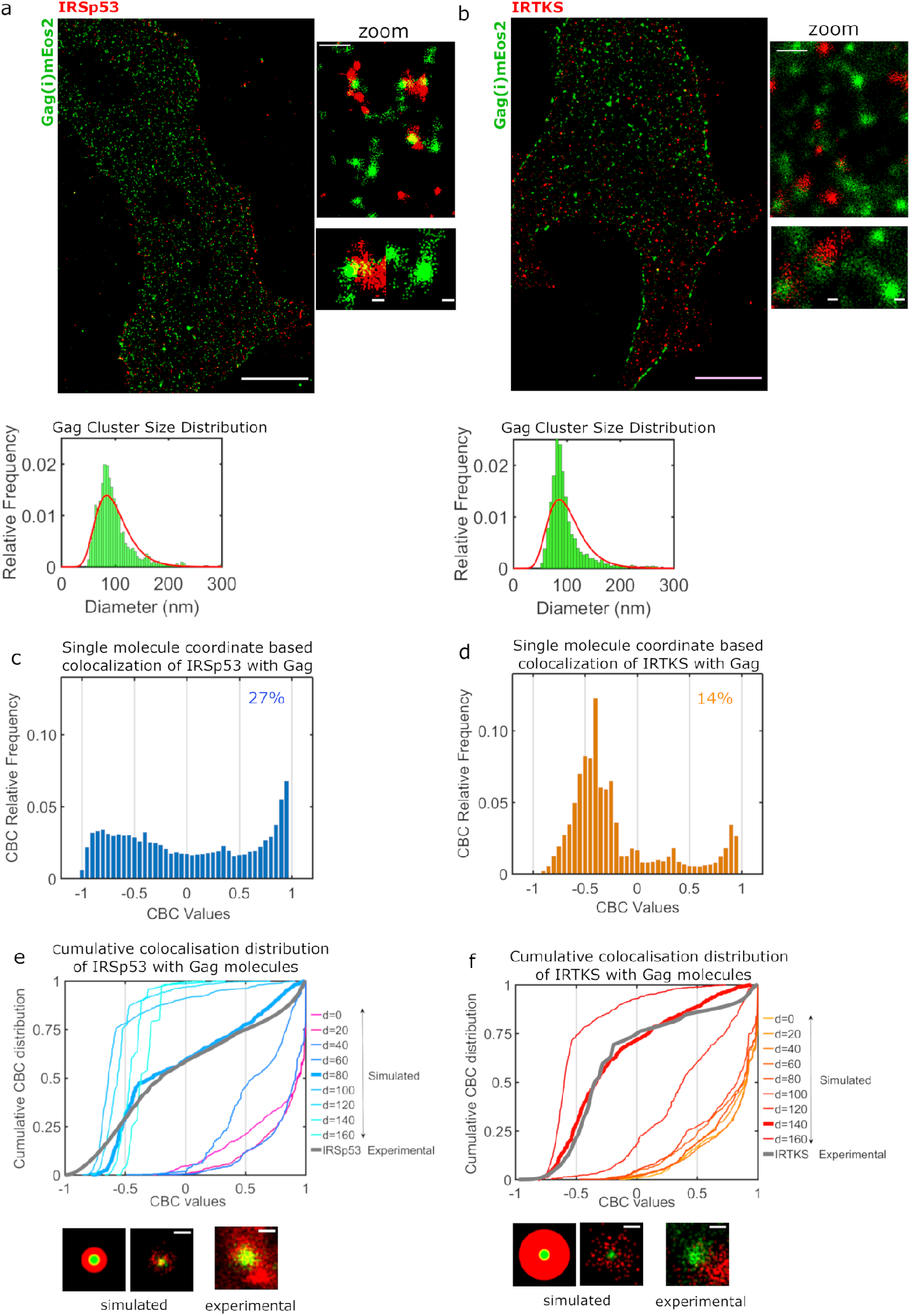
Super resolution microscopy imaging and simulations reveal preferential IRSp53 localization at HIV-1 Gag budding sites. a) Super-resolved dual color images of HEK293T cells expressing Gag(i)mEos2 immuno-labelled for IRSp53 (scale bar 10 μm) with magnified view (scale bar 500 nm) and Gag cluster size distribution. HIV-1 Gag assembly sites at the plasma membrane coincide strongly with cellular structures marked by IRSp53 (left panel, and magnified views) using coupled PALM/STORM microscopy in TIRF mode. HIV-1 Gag molecules localization density based scan (DBScan) analysis reveals clusters size distribution within the known range of HIV-1 particles size (80-150nm). b) Super-resolved dual color images of cells expressing Gag(i)mEos2 immuno-labelled for IRTKS (scale bar 10 μm) with magnified view (scale bar 500 nm) and Gag cluster size distribution. c) Quantification of coordinate based colocalization (as in (Malkusch et al., 2012)) at Gag assembly sites: CBC values for IRSp53 (c) and IRTKS (d) were plotted as relative frequencies. c) IRSp53 CBC values show a peak highly correlated (>0.5) with 27% of IRSp53 localizations highly correlated with Gag (>0.5). d) IRTKS CBC values show a peak anti-correlated/non correlated (−0.5 to 0). d) Comparison of cumulative frequency distributions of IRSp53/Gag and IRTKS/Gag. Only 14% of IRTKS localisations are highly correlated, with a peak of negatively correlated values. e) Comparison of experimental IRSp53/Gag CBC cumulative distribution with the simulated values. (See Fig S5 for details). IRSp53 shows an experimental cumulative CBC distribution corresponding to a belt of 80nm width (waist=40nm) centered at a distance of 80 nm from the center of Gag assembly sites (left graph, bold grey line corresponds to experimental data for IRSp53, bold blue line corresponds to simulated values closest to experimental data). IRSp53 thus corresponds to restricted pattern in and around a Gag assembly site (panel 1 schematic of simulated data, panel 2 simulated data and panel 3 experimental data). f) IRTKS experimental CBC distribution (bold grey line in graph) correspond to simulated ones using a belt of 200nm width (waist=100nm) centered at a distance of 140nm from the center of Gag assembly site (bold red line). IRTKS belt surrounding assembly sites is more diffuse and spreads out (panel 1 schematic of simulated data, panel 2 simulated data and panel 3 experimental data). Scale bars in the panels are 100 nm.

### IRSp53 is incorporated in HIV-1 particles

To assess IRSp53 incorporation into HIV-1 Gag particles, we purified Gag virus-like particles (VLP) from cells transfected with Gag-mCherry and several GFP-tagged I-BAR domain proteins (Fig.4a). IRTKS displays functional redundancy with IRSp53 (Chou et al., 2017; Millard et al., 2007), being able to curve membranes. IRSp53-I-BAR-GFP construct only contains the membrane curving I-BAR domain of IRSp53. PH-PLCδ-GFP, a PI(4,5)P_2_ binding protein, was used as a control, because it binds PI(4,5)P_2_ but does not generate membrane curvature. Fluorescent VLPs purified from these transfected cells were then visualized for two colors (green: GFP and red: mCherry), and Mander’s coefficient was calculated as an indicator of incorporation of the ectopic (green) GFP-tagged proteins within the (red) Gag-mCherry VLPs (see Materials and methods) (Fig.4a). We found that correlation for IRSp53-GFP and Gag-mCherry was 0.95-1 (Fig.3b, Left Graph, Red Column), indicating that almost all Gag-mCherry VLPs contained IRSp53-GFP. When using the IRSp53-I-BAR domain alone, we also obtained a high Mander’s coefficient, ~0.8 (Fig.4b). In contrast, for IRTKS-GFP, the Mander’s coefficient was 0.4-0.5, indicating no significant correlation between IRTKS-GFP and Gag-mCherry. Taken together, these results show the preferential incorporation of IRSp53-GFP, or its I-BAR-GFP domain, into released HIV-1 Gag particles. To study the incorporation of endogenous IRSp53 in HIV-1 particles, cells were transfected with plasmids expressing either wild-type infectious HIV-1 or codon optimized immature HIV-1 Gag protein (without genomic RNA). The virus particles were purified through a 20%-sucrose cushion or further through continuous iodixanol density gradient (as in (39, 40)). IRSp53 was found to be associated with the viral particles in both conditions, i.e. in infectious HIV-1 and in Gag VLPs, indicating that Gag alone is sufficient to recruit IRSp53 in the viral particles (Fig.4c,d, “IRSp53”). Tsg101 also showed an association with viral particles in both conditions (Fig.4c,d, “Tsg101”), as reported previously (33,34, 38). In contrast, IRTKS was not associated neither with Gag-VLP nor HIV-1 particles (Fig 4c, d, “IRTKS”). Upon further purification (Fig.4e), IRSp53, and the ESCRT proteins, Tsg101 and ALIX, were found associated within the same fractions containing the HIV-1 Gag viral particles, together with other well-known viral particle cofactors such as CD81, CD63 tetraspanins (39, 40). Thus, endogenous IRSp53 is most probably incorporated in HIV-1 particles in a Gag dependent manner.

**Fig. 4.**
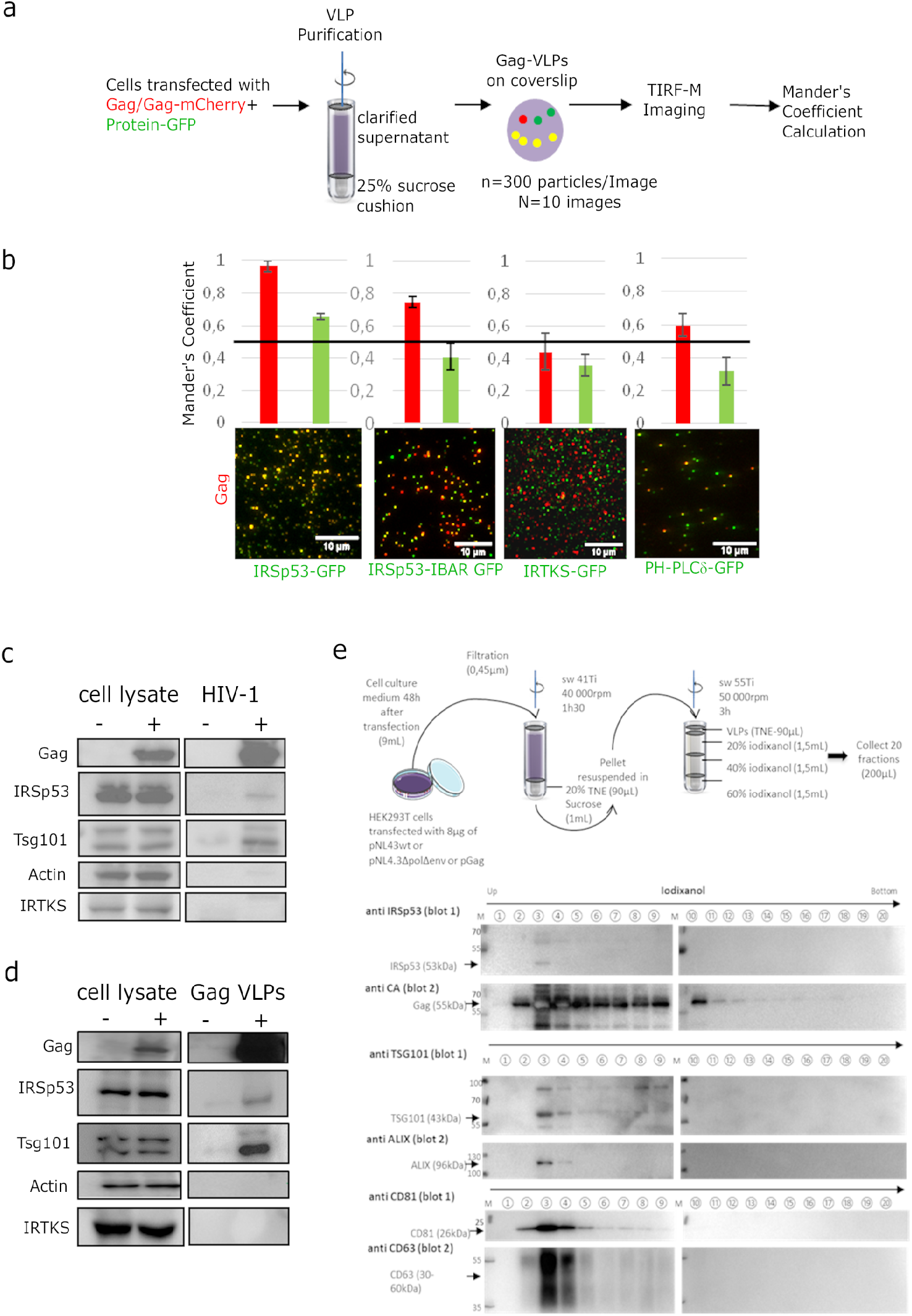
IRSp53 is incorporated into HIV-1 particles in a Gag dependent manner. a) Schematic for the protocol followed for imaging and analysis. HIV-1 Gag VLPs were purified from HEK293T cells expressing HIV-1 Gag/Gag-mCherry and IRSp53-GFP or other GFP tagged proteins (IRSp53-IBAR-GFP, IRTKS-GFP and PH-PLC**δ**-GFP). Purified Gag VLPs were then spotted over a poly-lysine treated glass slide and imaged by TIRF-Microscopy (particles were imaged in the red, and IRSp53 in the green channel). For each condition, 3000 particles (300 particles/image, 10 images) were counted. Fluorescence overlapping fraction (Mander’s coefficient, see Materials and Methods for details) were determined for Gag-mCherry and for IRSp53-GFP, IRTKS-GFP and PH-PLC**δ**-GFP and reported in the graphs. b) The 0,5 value indicates the random incorporation level (indicated by black line across the graph). IRSp53-GFP and IRSp53-IBAR show high correlation values (0.95-1 and 0.8, respectively). The other I-BAR domain proteins were not significantly correlated with Gag-mCherry particles (0.4-0.5). PH-PLC**δ**-GFP, a known marker of the phospholipid PI(4,5)P_2_, shows a slightly higher correlation (0.6), since HIV-1 Gag is known to associate with this phospholipid. c) Incorporation of IRSp53 into wild type pNL4-3 HIV-1 or d) Gag VLPs revealed by immunoblots against Gag(p24), IRSp53, IRTKS, Tsg101 or actin, as indicated. Following a 25% sucrose cushion purification, IRSp53 was found to be associated with released wild type HIV-1 (left panel) and Gag VLPs (right panel). Tsg101, known to be incorporated into released particles, was also found associated with viral particles. IRTKS, a closely related I-BAR protein to IRSp53, was not incorporated in purified HIV-1 viral particles or Gag-VLPs. e) Protocol of VLPs purification using sucrose cushions and iodixanol gradient. Briefly, pellets obtained after ultracentrifugation of cell culture medium of HEK293T transfected with pNL4.3HIV-1 or pGag were deposed on iodixanol gradient (20, 30 and 60%). 20 fractions of 200 μL were collected from top of the tube. Fractions collected following iodixanol gradient purification of NL4-3**Δ**Pol**Δ**Env Gag VLPs were analyzed using western blots for IRSp53 and Gag, TSG101 and ALIX, CD81 and CD63 revealed respectively on the same membrane (blot 1 and blot 2) revealing IRSp53 association with Gag viral particles and known cofactors.

### HIV-1 Gag is enriched at membrane tube tips generated by IRSp53 I-BAR domain

The results above demonstrate that not only IRSp53 is incorporated in Gag-VLPs, but it is present at the budding sites and its deletion strongly reduces HIV-1 particle release in a Gag-dependent manner by arresting its assembly at the cell plasma membrane (Fig.1). In order to advance molecular mechanistic understanding of the role of IRSp53 locally at Gag assembly sites, we assessed IRSp53 I-BAR/Gag interplay on model membranes, using GUVs. For Gag membrane binding assay, we first used a high concentration of IRSp53 I-BAR domain (0.5μM) while keeping PI(4,5)P2 concentration constant, in order to prevent having tubes generated by IRSp53 I-BAR (41) and to focus on analyzing Gag-flat GUV membrane binding efficiency in the presence of IRSp53 I-BAR. We found that Gag binding to GUV membranes is increased 7 fold when IRSp53-I-BAR domain was introduced first on GUVs before adding Gag (median value 6.7), compared to the condition of Gag only (median value 0.9) (p< 0.0001, Student’s t-test) (Fig.5a, “Gag only” vs. “I-BAR+Gag”). However, in the condition where Gag was introduced before adding the I-BAR domain, Gag intensity on GUV membranes increased only 3 fold as compared to the Gag only condition (p< 0.0001, Student’s t-test) (Fig.5a, “Gag only” vs. “Gag+I-BAR”). Notably, by comparing I-BAR+Gag and Gag+I-BAR conditions, we observed a 2-fold higher Gag intensity on GUV membranes in the first condition (p< 0.0001, Student’s t-test) (Fig.5a, “I-BAR+Gag” vs. “Gag+I-BAR”). Taken together, these results show that IRSp53 I-BAR facilitates Gag membrane binding on GUV in favor of increasing Gag concentration locally. Given these results, and that IRSp53 is a membrane curving protein involved in the early stage of generation of cell protrusions (20, 42), we asked whether the local membrane deformation induced by IRSp53 could be a preferred location for HIV-1 Gag assembly. Previous in vitro studies showed that when placing IRSp53 I-BAR domain outside PIP2-containing GUVs, even at a relatively low bulk concentration (about 0.02 - 0.06 μM), I-BAR domain can deform GUV membranes, generating tubes towards the interior of the vesicles (27,30, 43). We thus incubated GUVs with I-BAR domain at low concentration (0.05μM), which allows for the generation of inward membrane tubes, followed by the addition of Gag (Fig5b, see Materials and Methods). This experiment revealed that Gag was sorted preferentially to the tips of the tubes generated by the I-BAR domain (Fig.5b,c, Fig.S11 c,d). Furthermore, we observed that addition of HIV-1 Gag resulted in the formation of shorter I-BAR tubules as compared to GUVs incubated with the IRSp53-I-BAR domain alone (Fig S11 c,d), indicating an interference in I-BAR tubule elongation when Gag sorted to the tubule tips, suggesting that Gag usurps the IRSp53 tubulation function. Together, these results demonstrate that HIV-1 Gag binding to membranes is enhanced locally by the presence of IRSp53 I-BAR domain, and that Gag preferentially binds to highly curved membranes generated by the I-BAR domain of IRSp53.

**Fig. 5.**
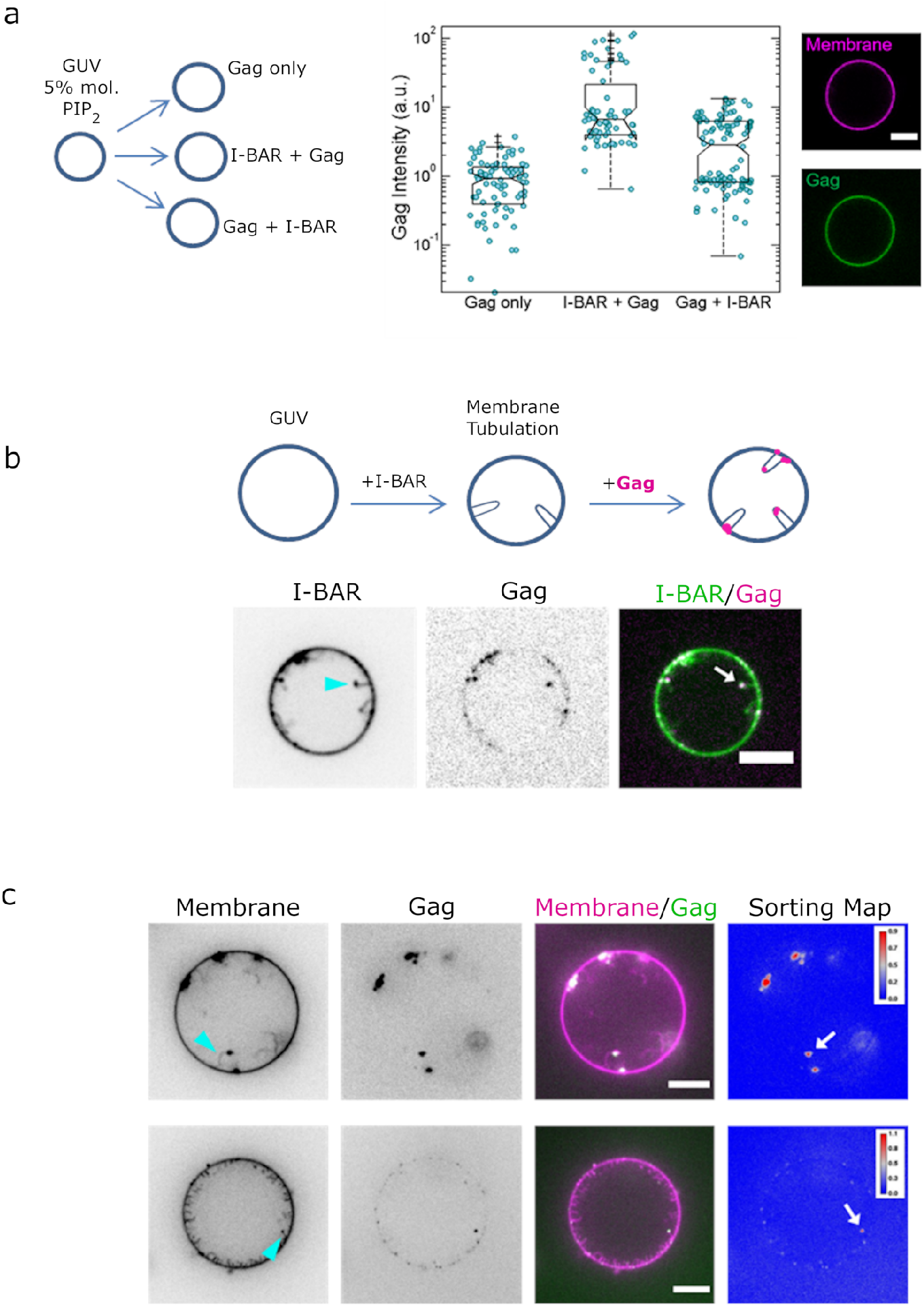
IRSp53 I-BAR domain enhances Gag recruitment to GUV-membranes and at the tip of I-BAR domain-induced tubes. a) (Left) AX488 Gag fluorescence intensity on membranes in the absence of I-BAR domain (named “Gag only”), in the presence of I-BAR domain where GUVs were first incubated with I-BAR domain and then Gag (named “I-BAR + Gag”) and GUVs were first incubated with Gag and then I-BAR domain (named “Gag + I-BAR”). Each circle presents one GUV analysis. N = 82 GUVs, n = 4 sample preparations for “Gag only”, N = 67 GUVs, n = 4 sample preparations for “I-BAR + Gag”, and N = 104 GUVs, n = 4 sample preparations for “Gag + I-BAR”. To pool all data points from the 4 sample preparations, in each preparation for all three conditions, Gag intensities were normalized by the mean Gag intensity in the “Gag only” condition. Protein bulk concentrations: 0.3 μM for AX488 Gag and 0.5 μM for I-BAR domain (not fluorescently labelled). (Right) Representative confocal images of AX488 Gag on GUV membranes in “I-BAR + Gag” condition. b) Representative confocal images of AX594 Gag in I-BAR domain-induced tubules. Inverted grayscale images are shown for I-BAR domain and Gag. Protein bulk concentrations: 0.3 μM for AX594 Gag and 0.05 μM for I-BAR domain (70% unlabeled and 30% AX488 labeled I-BAR domain). Cyan arrowhead points out a I-BAR domain-induced tubule and white arrow indicates the co-localization of Gag and I-BAR domain at the tip of the tubule. c) Representative confocal images of AX488 Gag in I-BAR domain-induced tubules. Inverted grayscale images were shown for membranes and Gag. Protein bulk concentrations: 0.3 μM for AX488 Gag and 0.05 μM for I-BAR domain (not fluorescently labelled). Cyan arrowheads point out I-BAR domain-induced tubules and white arrows indicate the co-localization of Gag and I-BAR domain at the tip of the tubules. Sorting map was obtained by calculating the fluorescence intensity ratio of Gag and membranes (see Material and Methods for more details). Scale bars, 5 μm.

## Discussion

The findings of this study uncovered the role of the host cellular I-BAR factor IRSp53 in HIV-1 Gag assembly and membrane curvature upon bud formation. In vitro, we showed that the IRSp53 I-BAR domain enhances Gag membrane binding locally (Fig5). Independently, we also checked if siRNA mediated knockdown of IRSp53 was changing total membrane binding of Gag using membrane flotation assays, but we did not observe any significant changes (not shown), emphasizing the local role of IRSp53 at Gag assembly sites rather than at the global cellular level. Indeed, IRSp53 was found at, or in the close vicinity of, Gag assembly platforms at the cell membrane (Fig.3), and is incorporated into Gag virus-like particles and in HIV-1 virions (Fig.4). Importantly, we revealed that IRSp53 partial knockdown arrests Gag assembly at the mid-bud formation stage (Fig.1) and that Gag preferentially locates at the tube tips induced by IRSp53 I-BAR domain, interfering with its long tubule formation in vitro (Fig.5). Altogether, IRSp53 appears instrumental in membrane curvature upon HIV-1 budding and is locally subverted as an essential factor needed for full HIV-1 Gag particle assembly. Using GUVs, we observed that Gag not only colocalizes with the IRSp53 I-BAR domain on the vesicles, but that the IRSp53-I-BAR domain increases Gag binding to these model membranes, mimicking the possible local Gag/IRSp53 interplay at the assembly site (Fig5a). Indeed, BAR domain proteins, in general, and IRSp53, in particular, are known to induce strong PI(4,5)P2 clusters (27, 44), and PI(4,5)P2 was shown to play a role in Gag binding to cell plasma membrane (45), as well as to be strongly clustered during virus assembly (9,28, 45). Thus, these results suggest that membrane binding of Gag on IRSp53-enriched membrane domains could promote the plasma membrane binding of both proteins (Fig.3, S6). This is in agreement with super resolution imaging in the cell, where Gag/IRSp53 interactions may take place at the Gag assembly sites, since, here, IRSp53 was localized in close proximity to Gag assembly sites in HEK293T cells (Fig.3) and in host CD4 T cells (Fig S9, S10). Our experiments suggest that Gag and IRSp53 are associated in a common complex at the cell plasma membrane (Fig2). The fact that upon Gag expression IRSp53 increases to the cell membrane (Fig.2b, c), suggests that Gag could activate IRSp53, through Rac1 activation (25), perhaps by releasing its auto-inhibition (24). This remains to be tested. HIV-1 particles are known to incorporate a large number of cellular proteins, many of which are directly involved in virus budding (38). Here, we showed that IRSp53 is incorporated in Gag-VLPs, as well as in purified HIV-1 virions (Fig.4), which most likely depends on the I-BAR domain of IRSp53 (Fig.4). Using the IRSp53-I-BAR domain on GUVs, we induced membrane protrusions that have a negative mean curvature similar to a viral bud; Gag was found enriched in these tubular structures but particularly at the tips that have a half-sphere geometry similar to a viral bud (Fig.5). This indicates that Gag binds preferentially to IRSp53 I-BAR-curved membranes in vitro in contrast to other almost-flat areas of the GUVs. Similarly, in cells, single molecule localisation images reveal some Gag clusters enriched at IRSp53 labelled protrusions at the plasma membrane (Fig S6a). IRSp53 clusters have already been reported prior to filopodia formation (20) and in negatively curved area at the onset of endocytic buds (42). Moreover, it was shown that inducing local membrane curvature helps to initiate Gag lattice formation (14). We thus propose that IRSp53 induces local membrane curvature, upon activation through Rac1/Cdc42 and effectors, which promotes local Gag recruitment and initiation of the viral assembly (knowing that expression of Gag can activate Rac1 (25)). Although the presence of RNA can facilitate the growth of the Gag network (46), favouring membrane bending due to the intrinsic curvature of assembling Gag hexamers, coarse grained simulations of HIV-1 Gag assembly showed that, above a certain threshold, this Gag self-assembly is unable to overcome the free energy penalty required to curve the membrane. Here, we observed that siRNA knockdown of IRSp53 induces a decrease in viral particle production and arrests the assembly at half completion (Fig.1). Since IRSp53 stabilizes curvature by scaffolding (30), another role of IRSp53 could be to lower this free energy barrier involved in the progression of the budding process beyond the half-sphere geometry, by stabilizing long enough the bud curvature. This, either directly by organizing linearly around the assembly site (43) and mechanically constricting the nascent bud, or indirectly with the help of actin polymerization. Interestingly, Ku et al., also observed that 60% of assembling particles exhibit a pause around the midway mark of the assembly process (47). This pause can provide a temporal window for IRSp53 to intervene in the progression of HIV-1 particle assembly as we propose here. Finally, ESCRT recruitment occurs at the end of virus assembly, after the membrane has been curved, forming a vesicle ready to bud (48, 49). Overexpression of a mutant of the ESCRT protein Tsg101 was previously shown to block HIV-1 budding at a late stage, arresting the budding with a characteristic bulb shaped phenotype indicative of a defect in the late stage of the bud scission (50), in contrast with our observations with the IRSp53 siRNA phenotype (Fig.1, Fig.S3). Consequently, this suggests that Gag-IRSp53 association is ESCRT independent (as shown in Fig. S4) and occurs at an earlier stage of virus assembly. Another study (51) showed that angiomotin, which acts as an adaptor protein for HIV-1 Gag and the ubiquitin ligase NEDD4L, functions in HIV-1 assembly prior to ESCRT-I recruitment. Interestingly, angiomotin also contains a BAR domain (52), but it is canonically involved in inducing positive curvature, as opposed to the negative curvature induced by I-BAR IRSp53. Thus, it is possible that angiomotin functions in another way, for example, at the viral bud neck which has both positive and negative curvatures by facilitating ESCRT recruitment. IRSp53 itself is a scaffold protein for cofactors of cortical actin signalling (16) and we have previously shown that a Rac1 signalling pathway, including IRSp53, is involved in HIV-1 particle production (25). Thus, it is possible that IRSp53 could also play a role in generating local cortical actin density in the vicinity of the viral bud. The role of cortical actin associated with IRSp53 scaffolding in that context remains to be elucidated. Our work illustrates a novel role for the host cellular I-BAR factor IRSp53, which is subverted by the retroviral Gag protein, in HIV-1 induced membrane curvature and in favoring the formation of the fully assembled viral particle.

## Material and Methods

### Antibodies

Rabbit polyclonal anti human IRSp53 (07-786 – Merck Millipore), mouse monoclonal anti CA (24.2 – NIH AIDS Reagent Program – FisherBioServices), mouse monoclonal anti GFP (B-2 – Santa Cruz biotechnology sc-9996). Rabbit polyclonal anti-human IRTKS (Bethyl), mouse monoclonal anti human CD63 (MX-49.129.5 – Santa Cruz biotechnology sc-5275), mouse monoclonal anti human CD81 (5A6 – Santa Cruz biotechnology sc-23962), rabbit polyclonal anti human IRSp53 (07-786 – Merck Millipore), mouse anti CA (183H125C – NIH3537), rabbit monoclonal anti human TSG101 (Abcam – ab125011), rabbit polyclonal anti human ALIX (Covalab – pab 0204).

### Plasmids

The plasmid expressing HIV-1 codon optimized Gag alone (pCMVGag [named pGag]), the plasmid expressing Pol and Env-deleted HIV-1 (named pNL4.3ΔPolΔEnv was a gift of E.Freed, HIVDRP, NIH, USA) encoding Gag alone with its packageable viral RNA (53) and the plasmid expressing full wild-type HIV-1 (named pNL4.3) were described previously (28). Plasmids IRSp53-GFP, IRTKS-GFP, PinkBAR-GFP and IRSp53-I-BAR-GFP were obtained from University of Helsinki (Finland) (27). Plasmids expressing PH-PLC*δ*-GFP was a gift of B.Beaumelle (IRIM, France), Gag(i)mCherry (named Gag-mCherry), Gag tagged with internal photo-activable mEos2 (named Gag(i)mEos2), p6-deleted Gag tagged with mEos2 (named pGagΔp6-mEos2) were described in (13).

### siRNA

Stealth siRNA (Invitrogen) targeting IRSp53 (BA-IAP2) and IRTKS (BAIAP2L2), and Smartpools (Dharmacon) targeting IRSp53 (BAIAP2) or random sequence for siRNA controls were used in this study.

### Cell culture and transfection

A human embryonic kidney cell line (HEK-293T) were maintained in Dulbecco’s Modified Eagle’s Medium (DMEM, GIBCO) and Jurkat T-lymphocytes were maintained in RPMI (GIBCO). Media were supplemented with 10% fetal bovine serum (FBS, Dominique Dutscher)), complemented with sodium pyruvate and antibiotics (penicillin-streptomycin) at 37°C with 5% CO2 atmosphere. HEK-293T cells were transfected as in (54). Based on different plasmids conditions, cells were transfected as follows (2*10^6^ cells/transfection): PLASMID, 8μg;. the amount of transfected plasmid was normalized by adding pcDNA3.1 empty plasmid. The cell medium was replaced 6 hours post-transfection and experiments were performed 24-48h post transfection. SiRNA transfections in HEK293T cells were performed with either RNAiMax (Invitrogen) or with JetPRIME (Polyplus) or by electroporation for Jurkat T cells. One day prior to transfection, 2×10^5^ cells/well were seeded in 2mL of growth medium without antibiotics, in a 6-well plate. Transfection was performed using the manufacturer’s protocol. 24 hours after siRNA transfection, the cells/well were again transfected using CaCl2/HBS. These cells were incubated at 37°C with 5% CO2 atmosphere, for 24/48 hours.

### Immunoprecipitation assay

HEK-293T cells were transfected following the calcium-phosphate technique. Based on different conditions plasmids were transfected as follows (2*10^6^ cells/transfection): Gag/pNLΔPolΔEnv (8μg each) and the amount of transfected plasmid was normalized by adding pcdna3.1 empty plasmid. The cell medium was replaced 6 hours post-transfection. 24 hours post-transfection, the cells were washed with cold 1X-PBS prior to collection with 800μL of chilled lysis buffer (50mM TRIS-HCl [pH=7.4]; 150mM NaCl; 1mM EDTA; 1mM CaCl2; 1mM MgCl2; 1% Triton, 0.5% sodium deoxycholate; protease inhibitor cocktail [Roche] 1 tablet/10mL lysis buffer). The cells were incubated on ice for 30 minutes and then centrifuged at 13,000 rpm/ 15minutes/ 4° C. The supernatant was collected in a new tube and the pellet was discarded. For each condition, 1000μg of protein (the collected supernatant) was incubated with 1μg of anti-IRSp53 antibody for overnight on a tube rotator at 4° C. 25μL of beads (Dynabeads Protein A, Life Technologies) was added to each tube of protein-antibody complex and incubated for 2 hours on the tube rotator at 4° C. The samples were then washed 5 times with the lysis buffer, followed by addition of 20μL 2X Laemmli’s buffer to the beads. The samples were denatured at 95° C for 10 minutes and then processed for Western blot.

### Western blot and analysis

50μg of each protein (intracellular) samples or 20μL of purified VLP samples, added with SDS loading dye, were resolved on a 10% SDS-PAGE gel. The gels were then transferred on to PVDF membranes. Immunoblotting was performed by incubating the membranes overnight with primary antibody at 4° C, and 2 hours with HRP conjugated secondary antibody at room temperature. The blot signals were detected using ECL Prime/ECL Select substrate (Amersham) and images were taken using Chemi-Doc (BioRad).

### VLP purification and quantification

24 or 48 hours post-transfection, culture supernatants containing Gag-VLPs were collected, filtered through a 0.45μM filter, and centrifuged first at 2000rpm/5mins/4° C and then at 5000rpm/5mins/4° C. The supernatant was then purified by loading it on a cushion of 25% sucrose (in TNE buffer) and ultracentrifuged at 100000g for 100 minutes (SW41Ti, Beckman Coulter) at 4° C. The pellets were resuspended in TNE buffer at 4°C overnight. Gag-VLP release was estimated by performing anti-CAp24 immunoblot and by quantifying Gag signal in the blots using ImageJ software as described in (25). The calculation for Gag-VLP release is: 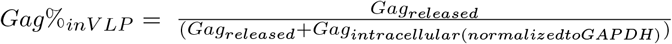.

### Membrane flotation assay

For each condition, 4×10^6^ cells were transfected and viral supernatants harvested 48h post-transfection, as described above. Cells were washed with ice-cold PBS and resuspended in Tris-HCl containing 4mM EDTA and 1X Complete protease inhibitor cocktail (Roche). Every step was then performed at 4° C. Cell suspensions were lysed using a dounce homogenizer, then centrifuged at 600g for 3min to obtain Post-Nuclear Supernatants (PNS). A cushion of 820μL of 75% (wt/vol) sucrose in TNE buffer (25mM Tris-HCl, 4mM EDTA, 150mM NaCl) was loaded at the bottom of an ultracentrifuge tube and mixed with 180 μL of PNS adjusted to 150mM NaCl. Two milliliters and 300 μ L of 50% (wt/ml) sucrose cushion followed by 0.9 mL of 10% (wt/ml) sucrose cushion were then layered to obtain the gradient that was then centrifuged in a Beckmann SW60Ti rotor at 35 000rpm, 4° C, overnight. Eight 500μL fractions were collected from the top to the bottom of the centrifuge tube and analyzed by western blotting.

### Electron microscopy

SiRNA treated HEK-293T cells were fixed in 4% paraformaldehyde and 1% glutaraldehyde in 0.1M phosphate buffer (pH 7.2) for 48h, washed with PBS, post-fixed in 1% osmium tetroxide for 1h and dehydrated in a graded series of ethanol solutions. Cell pellets were embedded in EPONŮ resin (Sigma) that was allowed to polymerize at 60°C for 48h. Ultrathin sections were cut, stained with 5% uranyl acetate and 5% lead citrate and deposited on colloidon-coated EM grids for examination using a JEOL 1230 transmission electron microscope.

### Sample preparation for super resolution PALM/STORM microscopy

HEK293T cells expressing HIV-1 Gag/Gag(i)mEos2 cultured on poly-l-lysine (Sigma) coated 25 mm round *♯* 1.5 coverslips (VWR) were fixed using 4%PFA + 4% sucrose in PBS for 15 min at RT. Samples were subsequently quenched in 50mM NH4Cl for 5 min. Samples were then washed in dPBS and then blocked for 15 min in room temperature using 1%BSA in PBS and subsequently in 0.05% Saponin in 1%BSA in PBS. Samples were stained using a 1:100 dilution of the primary antibodies (rabbit polyclonal anti-human IRSp53, Sigma and rabbit polyclonal anti-human IRTKS antibody, Bethyl) for 60 min in room temperature. Samples were washed 3×5 min using 1% BSA in PBS followed by 60 min staining using a 1:2000 dilution of the anti-rabbit Atto647N antibody (Sigma). Samples were washed 3×5 min with PBS and stored in light protected container in +4° C until imaged. Samples were mounted on a StarFrost slide with a silicon joint with the STORM buffer (Abbelight). Cells were imaged within 60 minutes after application of buffer.

### PALM/STORM Imaging

Single-molecule localization microscopy was performed on a Nikon inverted microscope equipped with 405-, 488-, 561- and 642-nm lasers, an EM-CCD Evolve 512 Photometrics camera (512*512, 16μm pixel size) with an oil immersion objective 100X NA1.49 Plan Apochromat. PALM imaging of Gag mEos2, activation was performed with lasers irradiance set to 0.3 kW/cm^2^ for 405 nm conversation and ~2.2 kW/cm^2^ for 561 nm excitation. Illumination was performed over a 25×25 μm^2^ area in the sample (1/e^2^ spatial irradiance distance) in TIRF-mode. 20-50,000 images were acquired for each cell with 50 ms integration time. The mean precision localisation in PALM measurements was found to be 20±5 nm (Fig S3). 2DSTORM imaging of Alexa647, was performed using a ~5 kW/cm^2^ irradiance with the 642 nm excitation. 25000 images were acquired for each condition. Image reconstruction was performed using the ThunderStorm plugin of ImageJ using Tetraspeck 100 nm multicolor beads (Life Technologies) as fiducial markers to correct for drift.

### Super Resolution Microscopy Analysis

The module DBSCAN of the super resolution quantification software SR Tesseler (55) was used to analyse the PALM localizations for quantification of Gag cluster sizes. In order to monitor the localisation of I-BAR proteins in the vicinity of Gag assembling particles, a binary mask was introduced into the PALM images. The centre of each Gag assembling cluster was determined and a custom MATLAB (Mathworks) code was used to extract localizations in a radius of 80nm around each Gag cluster center and to extract IBAR proteins localisations belonging to a disk of 150nm radius around the center of each Gag clusters. These subsets of coordinates were then used to calculate coordinate-based colocalization (CBC), developed by Malkusch *et al*. (37), and implemented in the ThunderSTORM plugin of ImageJ (Fig. S4). The coordinate-based colocalization (CBC) values are calculated from single-molecule localization data of two species (Gag and IBAR proteins (IRSp53 or IRTKS)). A CBC value is assigned to each single localization of each species. We analyzed the distributions of these CBC values by plotting and comparing the cumulative frequency distributions of the CBCs obtained in the two conditions (IRSp53 vs IRTKS). Finally, we performed a set of numerical simulations of images in ImageJ (see Fig. S8 legend for details) the average localization pattern of the immuno-labelled I-BAR proteins with respect to the mEos2 tagged Gag in PALM (Fig S5).

### Preparation and Imaging of Fluorescent VLPs

24h after seeding 2.10^6^ HEK293T cells were transfected with 8μg of pI-BAR-GFP proteins with or without 8μg of pGag/pGag(i)mCherry (2/3 1/3 respectively). 24h after transfection cells media (9mL) were filtered before performing VLPs purification by ultracentrifugation (SW41Ti rotor (Beckman) 29 000rpm, 1h30) on TNE 20% sucrose cushion. Pellets were resuspended with 110μL of TNE and allowed to sediment on round 25mm coverslips during 45 minutes in a chamber. VLPs were imaged with a Nikon Ti Eclipse 2 TIRF microscope. Images were taken with an Evolve EM-CCD camera – 512 photometrics, using a NA=1.45, 100X objective and 488 and 561nm lasers.

### Image Analysis for Colocalization

Images were acquired with Zeiss LSM780 (for fixed cells) or Nikon Eclipse Ti-2 in TIRF mode (for fluorescent viral particles). Colocalization analysis based on Mander’s coefficients was performed using JaCOP (Just another Colocalization Plugin) (56). Mander’s coefficient are defined as 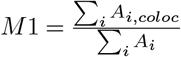 and 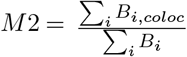, A and B being the two respective channels (mCherry and GFP). 0<M<1, with 1 full colocalization and 0.5 random colocalization. The M1 and M2 coefficients were calculated for several images and then represented as column graphs with red columns representing the degree of overlap of mCherry images with GFP images, and green columns representing the inverse. Iodixanol gradient. Cell culture medium of HEK293T (2.5×10^6^ cells plated) transfected with 8μg of pNL4.3ΔpolΔenv were collected 48h after transfection and filtered by 0.4*mw*m filter. The medium was then ultracentrifuged on a TNE 20% sucrose cushion using a SW41Ti rotor (Beckman) at 40,000rpm during 1h30. A solution with 0.25M sucrose, 1mM EDTA, 10mM tris HCL pH 7,4 was used to diluted the 60% iodixanol stock solution (OptiPrep from Sigma) and to prepare 40% and 20% iodixanol solution. 1,5mL of each dilution (60%, 40% and 20% iodixanol) were successively layered in a SW55Ti tube (Beckman) and the pellet of VLPs obtained after ultracentrifugation on sucrose cushion was loaded on the top. The tube was ultracentrifuged at 50 000rpm with a SW55Ti rotor (Beckman) during 3h. Then 20 fractions of 200μl were collected from top of the tube. 20μL or 18.75μL of each fraction were loaded for western blotting.

### GUV reagents

Brain total lipid extract (131101P) and brain L-α-phosphatidylinositol-4,5-bisphosphate (PIP2, 840046P) were purchased from Avanti Polar Lipids/Interchim. BODIPY-TR-C5-ceramide, (BODIPY TR ceramide, D7540) and Alexa Fluor 488 C5-Maleimide (AX 488) were purchased from Invitrogen. β-casein from bovine milk (>98% pure, C6905) and other reagents were purchased from Sigma-Aldrich. Culture-Inserts 2 Well for self-insertion were purchased from Ibidi (Silicon open chambers, 80209).

### Protein purification and fluorescent labelling

Recom-binant mouse IRSp53 I-BAR domain was purified and labeled with AX488 dyes as previously described (Prévost et al., 2015; Saarikangas et al., 2009). Recombinant HIV-1 immature Gag protein was purified by J. Mak as described in (Yandrapalli et al., 2016) and labelled with Alexa488 maleimide dyes (Invitrogen). Briefly, a 200μM solution of the maleimide dye was incubated overnight at 4°C with a 20μM solution of the Gag purified protein in a buffer of pH 8.0 with 1M NaCl and 50mM Tris. Post incubation, the labelled mixture was subjected to dialysis with the Slide-A-Lyzer Mini Dialysis Device (Thermo Scientific), following the manufacturer’s instructions to remove the excess unbound dye from the solution.

### GUV preparation and observation

#### Lipid and buffer compositions

Lipid compositions for GUVs were brain total lipid extract (Yu Seong-hyun et al., 2006) supplemented with 5 mole% brain PIP2. If needed, 0.5 mole% BODIPY TR ceramide was present in the lipid mixture as membrane reporter. The salt buffer inside GUVs, named I-buffer, was 50 mM NaCl, 20 mM sucrose and 20 mM Tris pH 7.5. The salt buffer outside GUVs, named O-buffer, was 60 mM NaCl and 20 mM Tris pH 7.5.

#### GUV preparation

GUVs was prepared by using the polyvinyl alcohol (PVA) gel-assisted method (Weinberger et al., 2013a). Briefly, a PVA solution (5% (w/w) of PVA in a 280 mM sucrose solution) was warmed up to 50°C before spreading on a coverslip that was cleaned in advance by being bath sonicated with 2% Hellmanex for at least 30 min, rinsed with MilliQ water, sonicated with 1M KOH, and finally sonicated with MilliQ water for 20 min. The PVA-coated coverslip was dried in an oven at 50°C for 30 min. 5-10 *mul* of the lipid mixture (1 mg/mL in chloroform) was spread on the PVA-coated coverslip, followed by drying under vacuum for 30 min at room temperature. The PVA-lipid-coated coverslip was then placed in a 10 cm cell culture dish and 0.5 mL −1 mL of the inner buffer was added on the coverslip, followed by keeping it stable for 45 min at room temperature to allow GUV to grow.

#### Sample preparation and observation

GUVs were first incubated with either Gag or I-BAR domain at bulk concentrations depending on the designed experiments for at least 15 min at room temperature before adding either I-BAR domain or Gag, respectively, into the GUV-protein mixture. In experiments where there was only Gag but no I-BAR domain, the stock solution of I-BAR domain was used in order to obtain a comparable salt strength outside GUVs as those where I-BAR domain was present. The GUV-protein mixture was then incubated at least 15 min at room temperature before observation. For Gag/I-BAR membrane recruitment assay, samples were observed by a Nikon C1 confocal microscope equipped with a 60X water immersion objective (Nikon, CFI Plan Apo IR 60X WI ON 1.27 WD 0.17). For Gag/I-BAR tubulation assay, samples were observed with an inverted spinning disk confocal microscope Nikon eclipse Ti-E, equipped with Yokogawa CSU-X1 confocal head, 100X CFI Plan Apo VC objective (Nikon) and a CMOS camera, Prime 95B (Photometrics). For all experiments, coverslips were passivated with a *β*-casein solution at a concentration of 5 g.L^-1^ for at least 5 min at room temperature. Experimental chambers were assembled by placing a silicon open chamber on a coverslip.

#### GUV Image analysis

Image analysis was performed by using Fiji (57).

#### Quantification of AX488 Gag binding on GUV membranes

Fluorescence images were taken at the equatorial planes of GUVs using identical confocal microscopy settings. The background intensity of the AX488 channel was obtained by manually drawing a line with a width of 10 pixels perpendicularly across the membrane of a GUV. We then obtained the background intensity profile of the line where the x-axis of the profile is the length of the line and the y-axis is the averaged pixel intensity along the width of the line. The background intensity was obtained by calculating the mean value of the sum of the first 10 intensity values and the last 10 intensity values of the background intensity profile. To obtain Gag fluorescence intensity on the membrane of the GUV, we used membrane fluorescence signals to find the contour of the GUV. Then, a 10 pixel wide band centred on the contour of the GUV was used to obtain the Gag intensity profile of the band where the x-axis of the profile is the length of the band and the y-axis is the averaged pixel intensity along the width of the band. Gag fluorescence intensity was then obtained by calculating the mean value of the intensity values of the Gag intensity profile, following by subtracting the background intensity.

#### Gag sorting map

Fluorescence images of GUVs were taken using identical confocal microscopy settings. For every GUV, we first calculated the fluorescence intensity ratio for every pixel of the Gag and membrane images of a GUV using 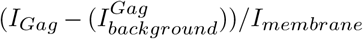, where *I_Gag_* is the Gag intensity, 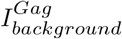 is the background intensity in the Gag channel, and *I_membrane_* is the membrane intensity. The sorting map was then obtained by converting the resulting image from the previous step to a pseudo-colored image via the “Look Up Table, Phase”. The background intensity value in the Gag channel was the man intensity value of a 50 pixels wide square in the background outside GUVs. The sorting map of I-BAR domain was obtained by using the same procedure as those for Gag.

#### Statistics

All notched boxes show the median (central line), the 25th and 75th percentiles (the bottom and top edges of the box), the most extreme data points the algorithm considers to be not outliers (the whiskers), and the outliers (crosses).

## ACKNOWLEDGEMENTS

The authors greatly acknowledge the Montpellier MRI-CNRS and CEMIPAI microscopy facility for access to the PALM/STORM microscopes. We thank Eric Freed (NIH, Frederick, MD USA) for providing the pNL43GagΔPolΔEnv plasmid and A.Cimarelli (CIRI, Lyon, France) for providing the pGag(myc) plasmid. The authors greatly acknowledge the Cell and Tissue Imaging (PICT-IBiSA), Institut Curie, member of the French National Research Infractucture France-Biolmaging (ANR10-INBS-04). DM and CF are members of the ImaBio Consortium of the CNRS (GDR ImaBio). This work was supported by the ANRS grant ECTZ35754. KI was the recipient of an ANRS fellowship for 3 years (2017-2020). RD is a recipient of a SIDACTION fellowship.

## Supplementary data

**Fig. S1.**
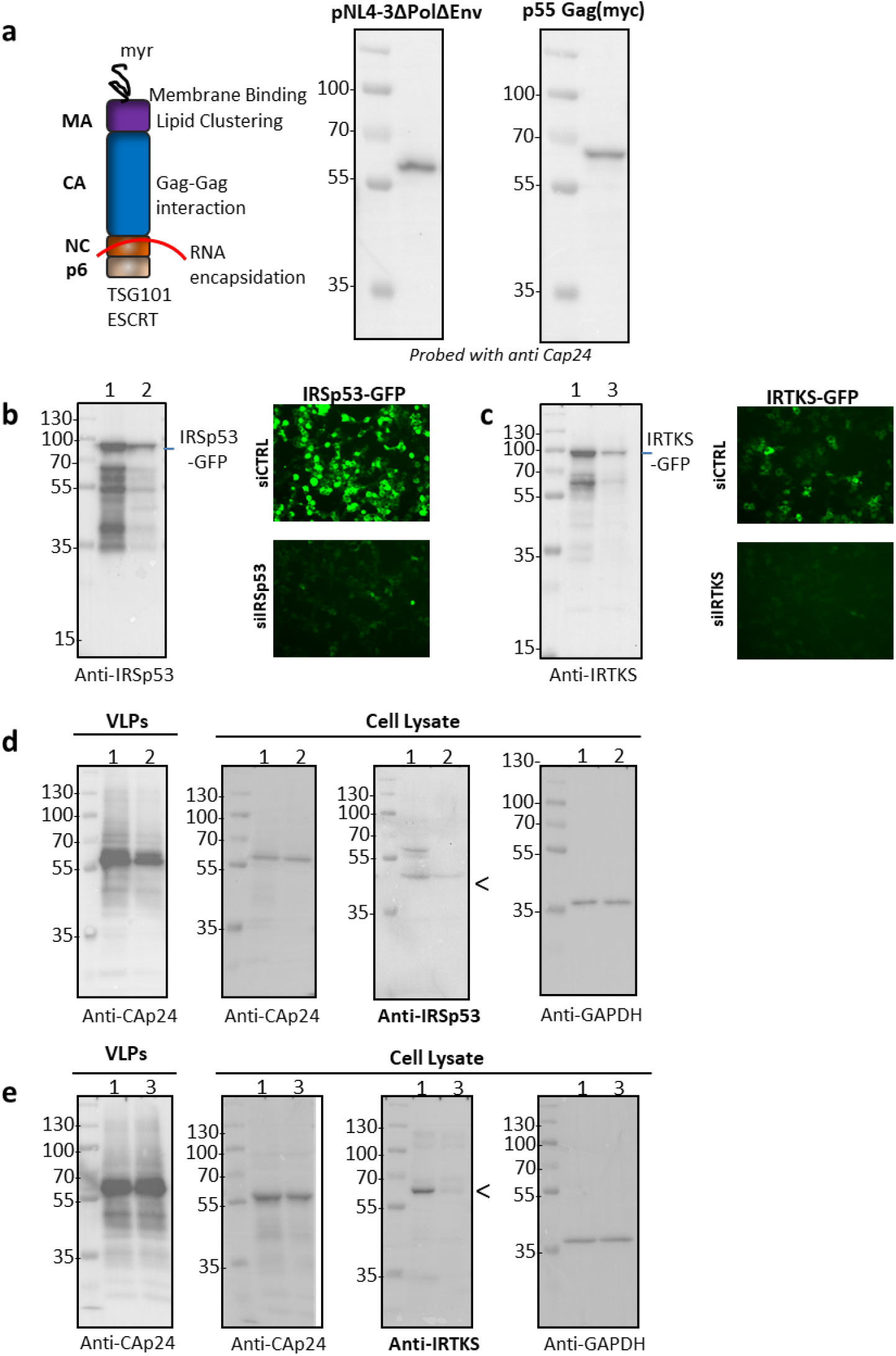
Effect of siRNA-based IRSp53 and IRTKS knockdown on HIV-1 Ga particle release. a) Expression of HIV-1 Gag in HEK293T cells (from molecular clones pNL43ΔPolΔEnv or pGag(myc)) and a schematic representation of the viral Gag structure. Gag is known to have four main domains, the matrix MA interacting with cell membrane, the capsid CA for Gag oligomerization, the nucleocapsid NC interacting with the viral genomic (+)RNA and p6 recruiting Tsg101 for particle budding. b) Validation of siRNA targeting IRSp53. siRNA targeting IRSp53 diminishes expression of IRSp53-GFP as seen by immunoblot (Left, Lane 1 siCTRL, Lane 2 silRSp53) and fluoresence imaging of 293THEK cells expressing IRSp53-GFP with corresponding siRNA (right). c) Validation of siRNA targeting IRTKS. siRNA targeting IRTKS diminishes expression of IRTKS-GFP as seen by immunoblot (Left), Left, Lane 1 siCTRL, Lane 3 siIRTKS) and fluorescence imaging of 293THEK cells expressing IRTKS-GFP with siRNA, as indicated (right). d) Reduced viral release are seen in cells knocked down for IRSp53 (representative immunoblots, Lane 1 siCTRL, Lane 2 siIRSp53). e) IRTKS knockdown has no significant effect on HIV-1 Gag particle release (representative immunoblots, Lane 1 siCTRL, Lane 3 silRTKS).

**Fig. S2.**
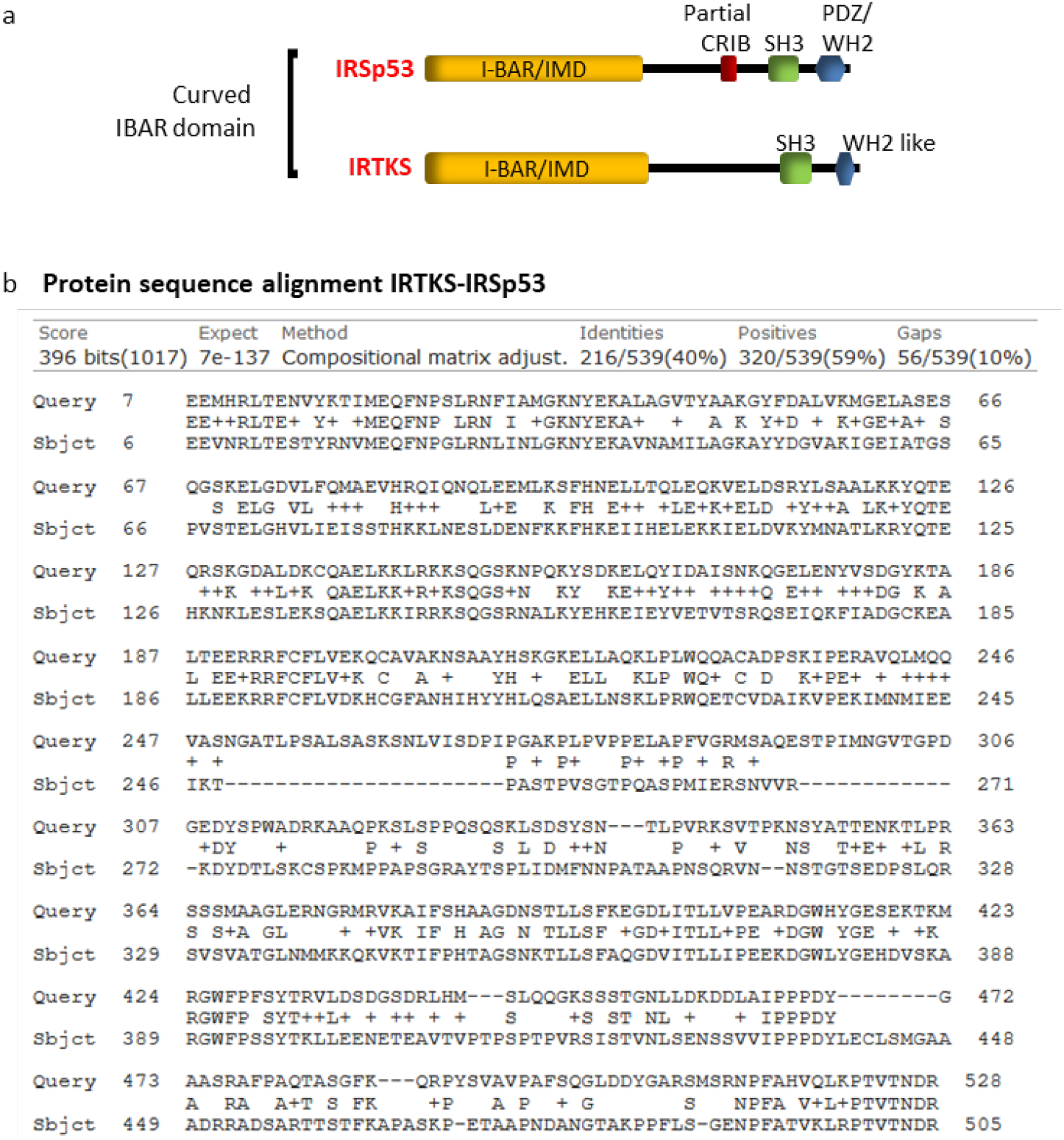
Protein sequence comparison of IRSp53 and IRTKS. a) IRSp53 and IRTKS are membrane curving I-BAR proteins. b) IRSp53 (Query, Accession Q9QB8) shares a 40% sequence homology with IRTKS (Subject, Accession Q9UHR4) Sequence alignment performed with NCBI Protein BLAST. Most of this homology is centered in the I-BAR/IMD domain of the three proteins and the C terminal SRC Homology 3 (SH3) domain common to the two proteins.

**Fig. S3.**
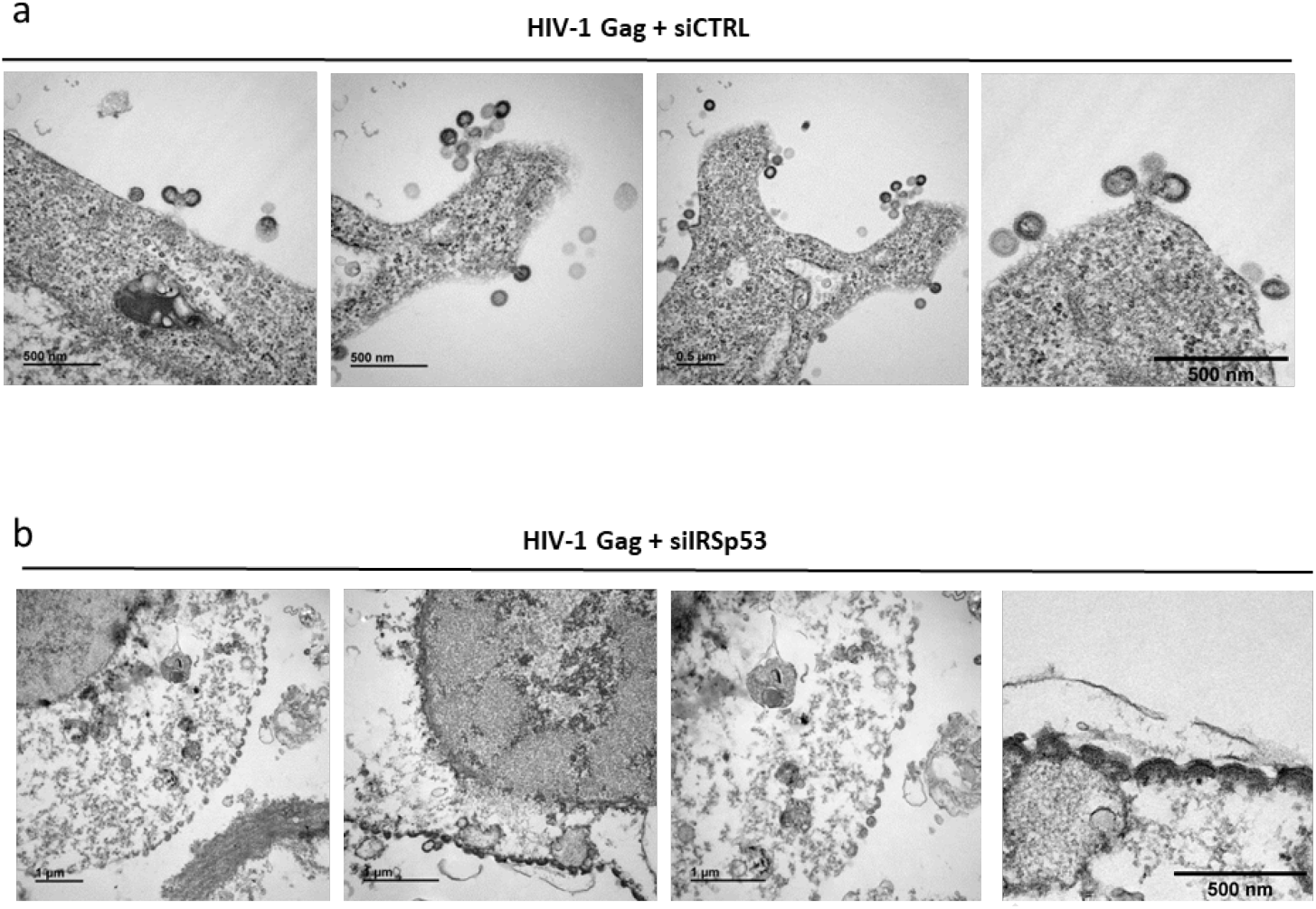
Transmission electron microscopy of siRNA treated HEK293T cells expressing HIV-1 Gag. **SiRNA control** (a) or siRNA IRSp53 (b) treated cells reveal that IRSp53 KD cells display HIV-1 Gag arrested assembly at the cell plasma membrane as shown by transmission electron microscopy.

**Fig. S4.**
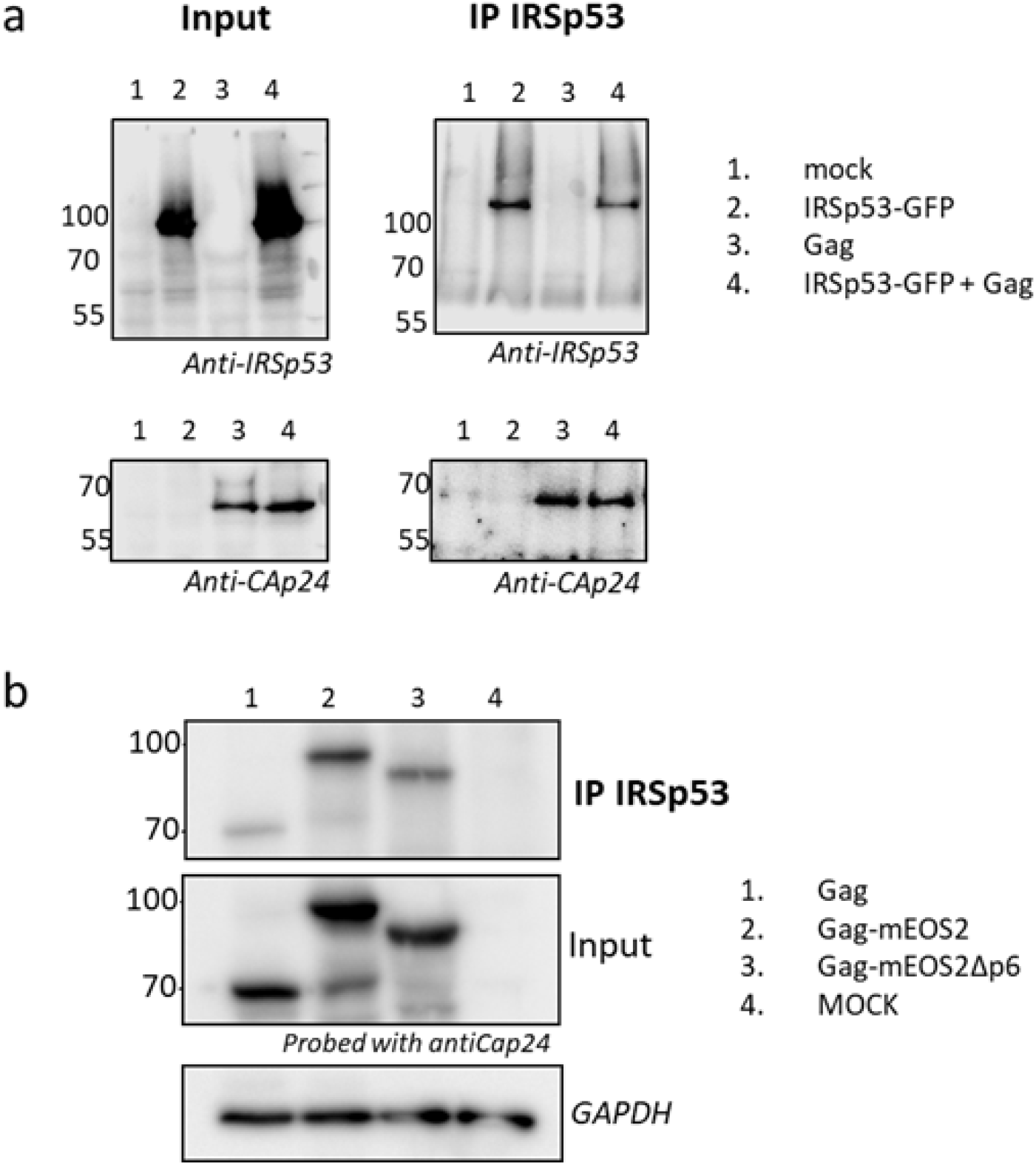
**a)Complexing of IRSp53-GFP with HIV-1 Gag**. Pulldown of IRSp53-GFP and IRSp53-GFP with Gag immuno-precipitated with an IRSp53 antibody (lane 2 and 4) and revealed with anti-CAp24 and anti-IRSp53 antibodies respectively using immunoblots. Mock (lane 1) and Gag (lane 3) are cell lysates from transfected HEK293T cells, used as controls. It is worth noticing that Gag (lane 3) is indeed immuno-precipitated (without ectopic IRSp53-GFP) due to the presence of endogenous IRSp53. **b) Complexing of IRSp53 with Gag-mEOS2 is independent of Gag-p6 domain**. Addition of an internal mEos2 tag within the Gag protein does not affect its complexing with IRSp53 (Compare lane 1 and Lane 2). Gag-mEos2Δp6, a mutant of Gag deficient in p6 dependent-ESCRT recruitment, is also immuno-precipitated (IP) with IRSp53 antibody (lane 3), indicating that loss of the p6 domain and Tsg101 recruitment is not a limiting factor for Gag-IRSp53 complexing. Lane 4 as an IP control.

**Fig. S5.**
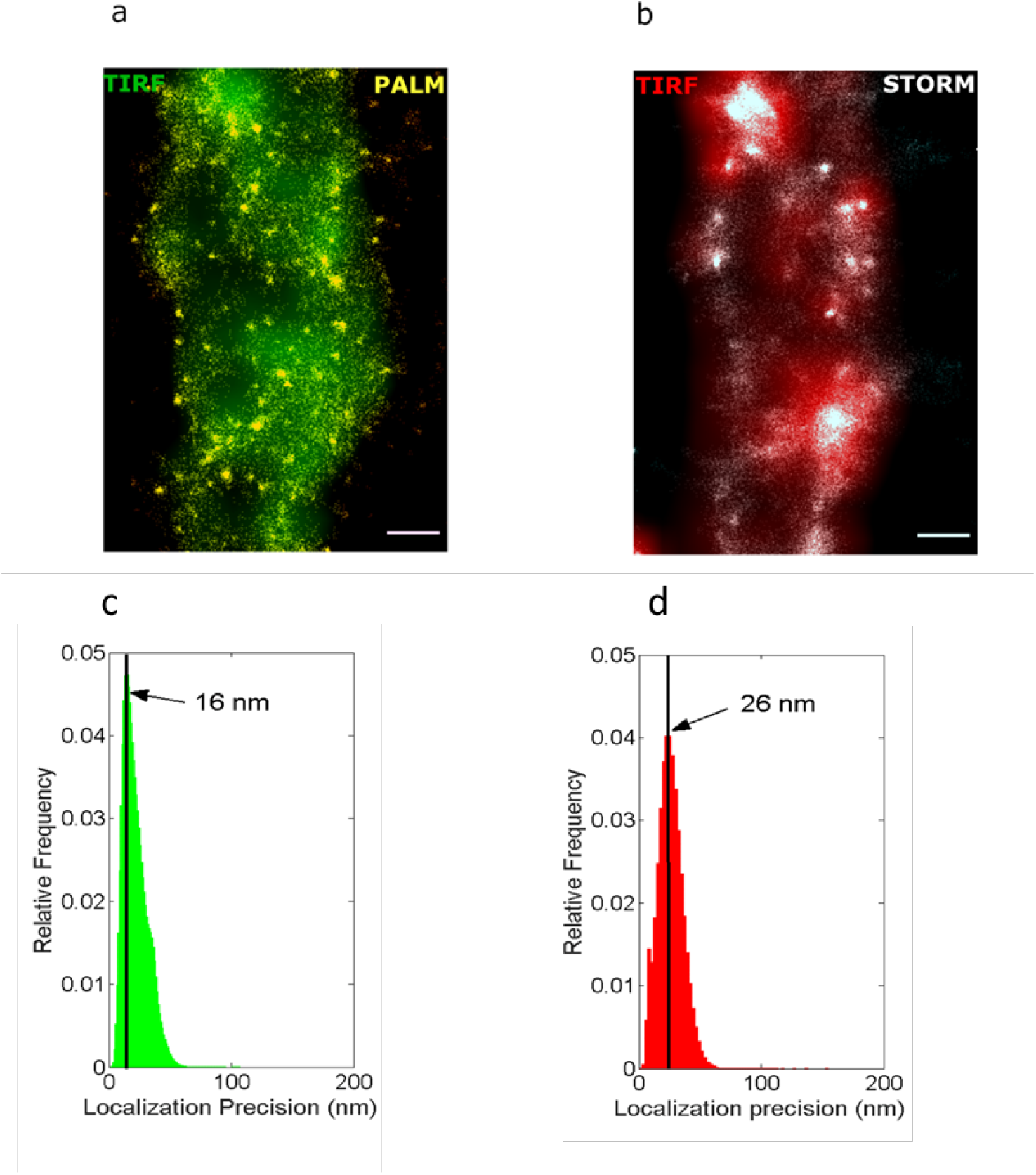
Super resolution PALM-STORM microscopy and localization precision. a) Single HIV-1 assembly sites detected using PALM (yellow) as compared to TIRF-M (green). b) Nanoscale structures detected in STORM (white) as compared to TIRF-M c) Localization precision in PALM and d) STORM measurements.

**Fig. S6.**
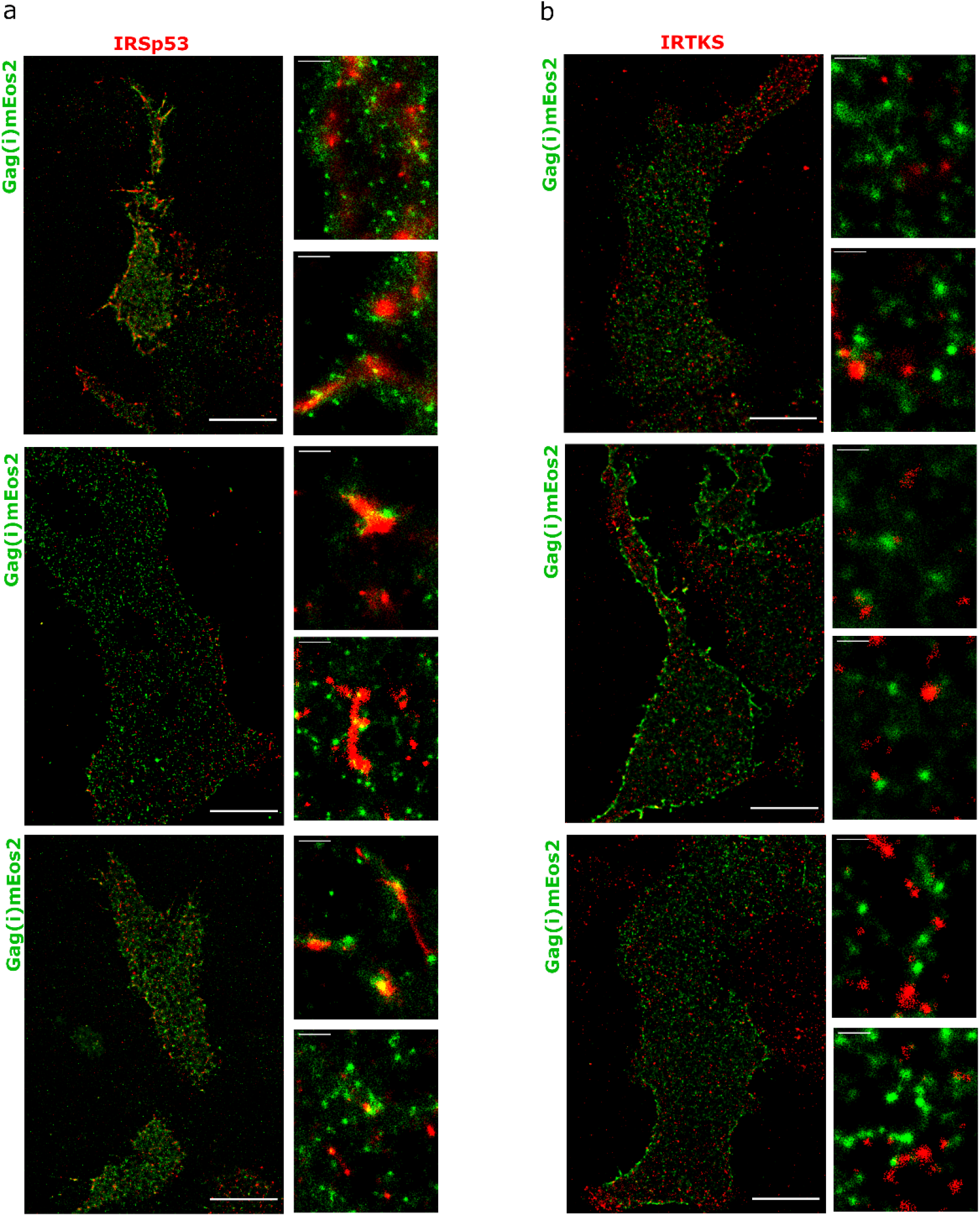
Super resolution PALM/STORM images of HEK293T cells expressing HIV-1 Gag(i)mEos2 (in green) and immunolabelled for a) IRSp53 (Atto647N, in red) and b) IRTKS (Atto647N, in red), with magnified images adjacent to the main image. Scale bar is 10μm for the large images and 500 nm for the magnified images.

**Fig. S7.**
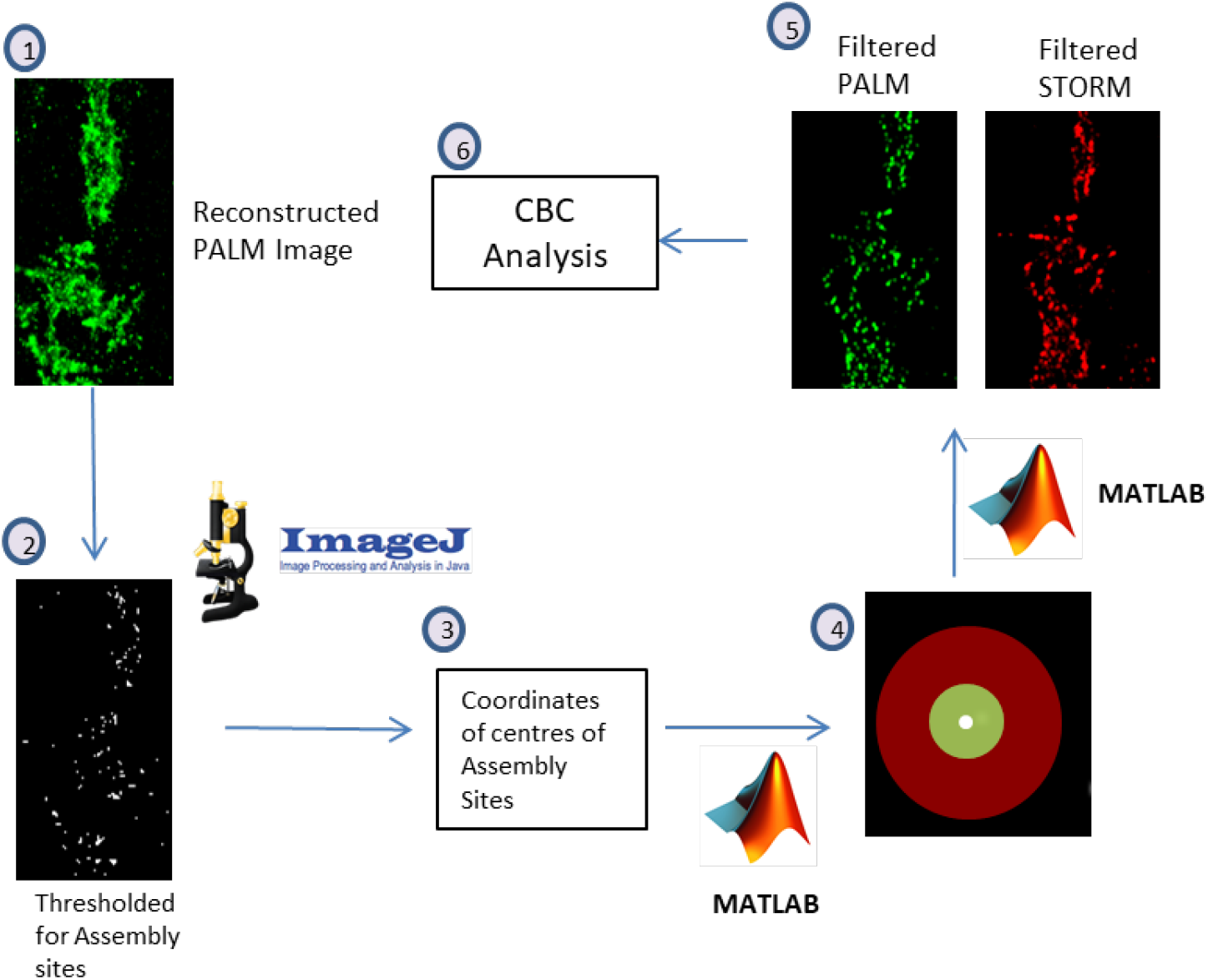
Workflow of image analysis for PALM/STORM images. (1) Reconstructed PALM images (2) The PALM images were thresholded in ImageJ to obtain a mask of the assembly sites of Gag. (3) The coordinates of centres of these assembly sites were then obtained using the Analyze Particles feature of ImageJ. (4) These coordinates were then used to analyze the original PALM and STORM single molecule localizations, and extract PALM coordinates from an area corresponding to size of assembly sites (r<80nm from center), and STORM coordinates in a larger area around the assembly site (r<150nm from center). (5) Using these filters, a new set of PALM and STORM localizations were generated, which corresponded to Gag assembly sites and the area around these sites. (6) These localizations were then used to calculate the coordinate based colocalization (CBC) for PALM (Gag) and STORM (IRSp53/IRTKS). The CBC values were then plotted as cumulative probability distributions.

**Fig. S8.**
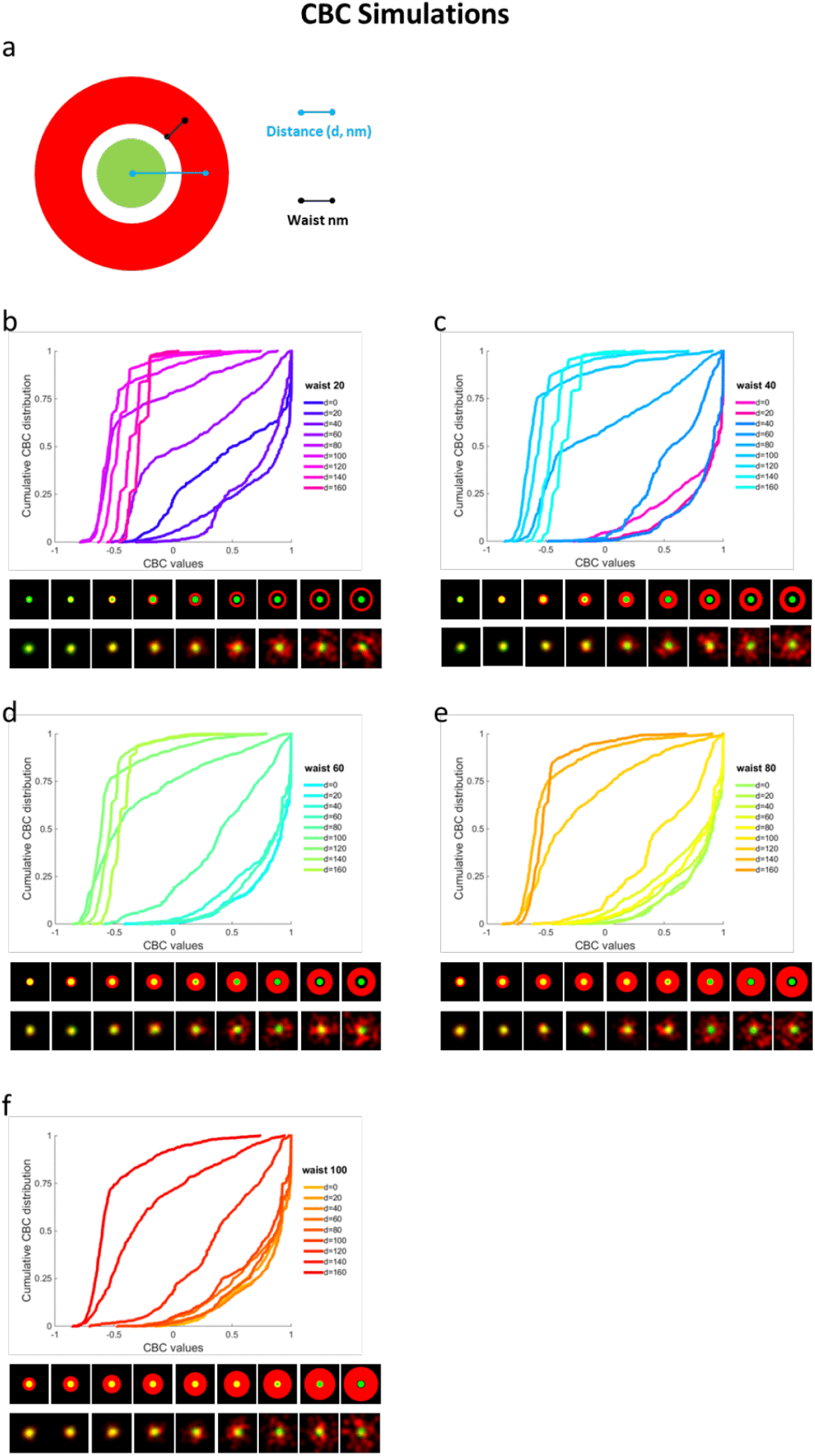
Cumulative frequency distributions of CBCs for simulated PALM/STORM data. Simulations corresponding to different patterns of STORM localizations around PALM clusters were performed using ImageJ. a) Each molecule position was randomly distributed within a fixed size disk of 100nm diameter for the PALM localizations (which represent 70% of all the experimental Gag cluster diameter found) or within a belt surrounding this disk for the STORM localization. The STORM belt size ranged from a waist of 20 nm (width 40nm) to a waist of 100 nm (width 200nm), with distances from center of the simulated viral bud (PALM disk) to the center of the surrounding belt ranging from 0 nm to 160 nm for each waist size. Each simulated localization was then convoluted by a 2D spatial Gaussian function with a waist of 200nm in x and y direction to simulate the point spread function of the microscope. The number of photons emitted by each simulated position was randomly distributed according a Gaussian distribution centered to a value equivalent to the one experimentally observed, this in order to obtain a localization precision equivalent to the experimental ones (see Fig S3 for the experimental localization precision). Each set of simulated images was then analyzed using the Thunderstorm plugin of ImageJ with the same parameter as the one used for experimental data. Single molecule localizations obtained from this analysis were then used for CBC analysis using the CBC plugin of Thunderstorm as it was done for experimental data sets (b to f).

**Fig. S9.**
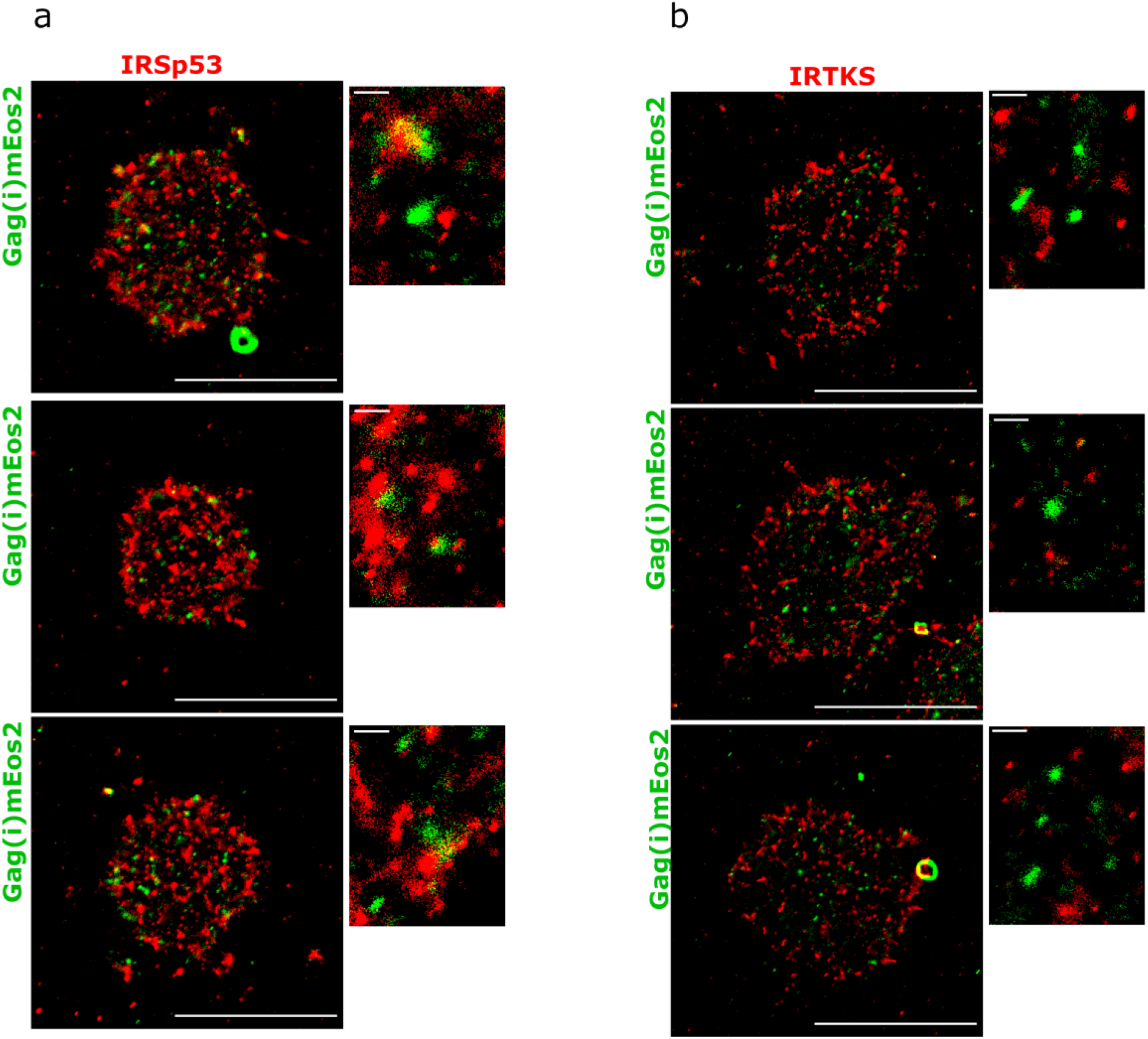
Super resolution PALM/STORM images of CD4+ Jurkat T cells expressing HIV-1 Gag(i)mEos2 (in green) and immunolabelled for a) IRSp53 (Atto647N, in red) and b) IRTKS (Atto647N, red), with magnified images adjacent to the main image. Scale bar 10μm for the large images and 500 nm for the magnified images.

**Fig. S10.**
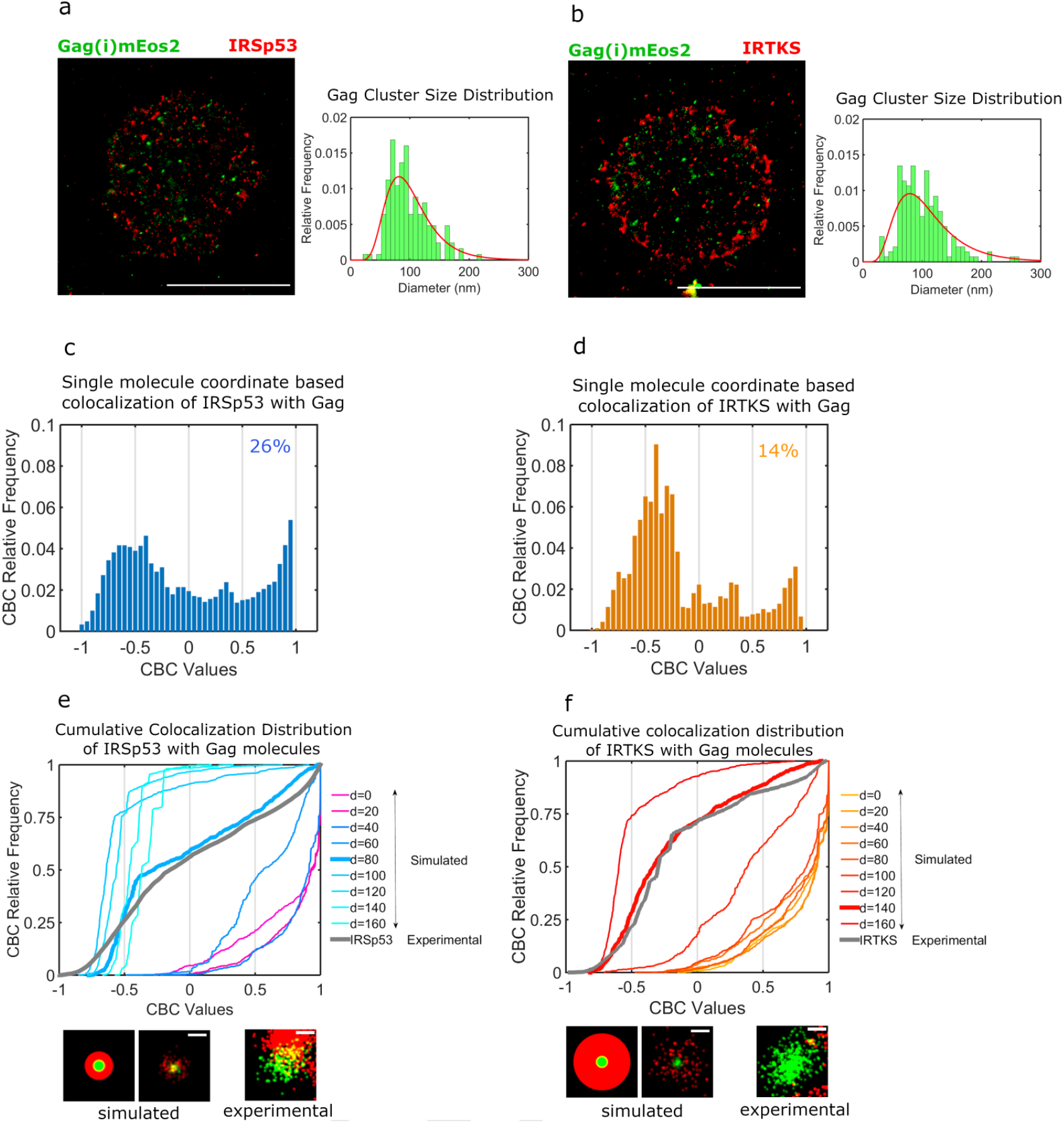
IRSp53, but not IRTKS, is localized at HIV-1 Gag budding sites in host CD4+ Jurkat T cells as revealed by super resolution microscopy. a, Super resolved dual color PALM/STORM images of Jurkat T cells immuno-labelled with IRSp53. As in Fig. 3, HIV-1 Gag molecules localization density based scan (DBScan) analysis reveals clusters size distribution within the known range of HIV-1 particles size (80-150nm). b, Jurkat cells immuno-labelled with IRTKS show a Gag cluster size distribution within the same range using the same DBScan analysis. CBC values for IRSp53 (c) and IRTKS (d) were plotted as relative frequencies. c, IRSp53 CBC values show a peak of IRSp53 localizations (26%) highly correlated with Gag (>0.5). d) IRTKS CBC values show a peak at anti-correlated and non correlated (−0.5 to 0). e) Experimental CBC values for IRSp53 and IRTKS were plotted as cumulative frequency distributions to be compared to simulated distributions obtained at different distances and structures (See Supplementary Fig.3). IRSp53 shows a cumulative CBC distribution corresponding to a simulation of waist at 40nm (width 80 nm) and a distance of 80nm (left graph, bold grey line corresponds to experimental data for IRSp53, bold blue line corresponds to simulated values closest to experimental data). IRSp53 thus corresponds to restricted pattern in and around a Gag assembly site (panel 1 schematic of simulated data, panel 2 simulated data and panel 3 experimental data). d, IRTKS experimental CBC distribution (bold grey line in graph) is close to simulations of waist 100nm (width 200nm) and a distance of 140nm (bold red line). IRTKS is more diffuse and spread out (panel 1 schematic of simulated data, panel 2 simulated data and panel 3 experimental data). Scale bar in the panels= 100nm.

**Fig. S11.**
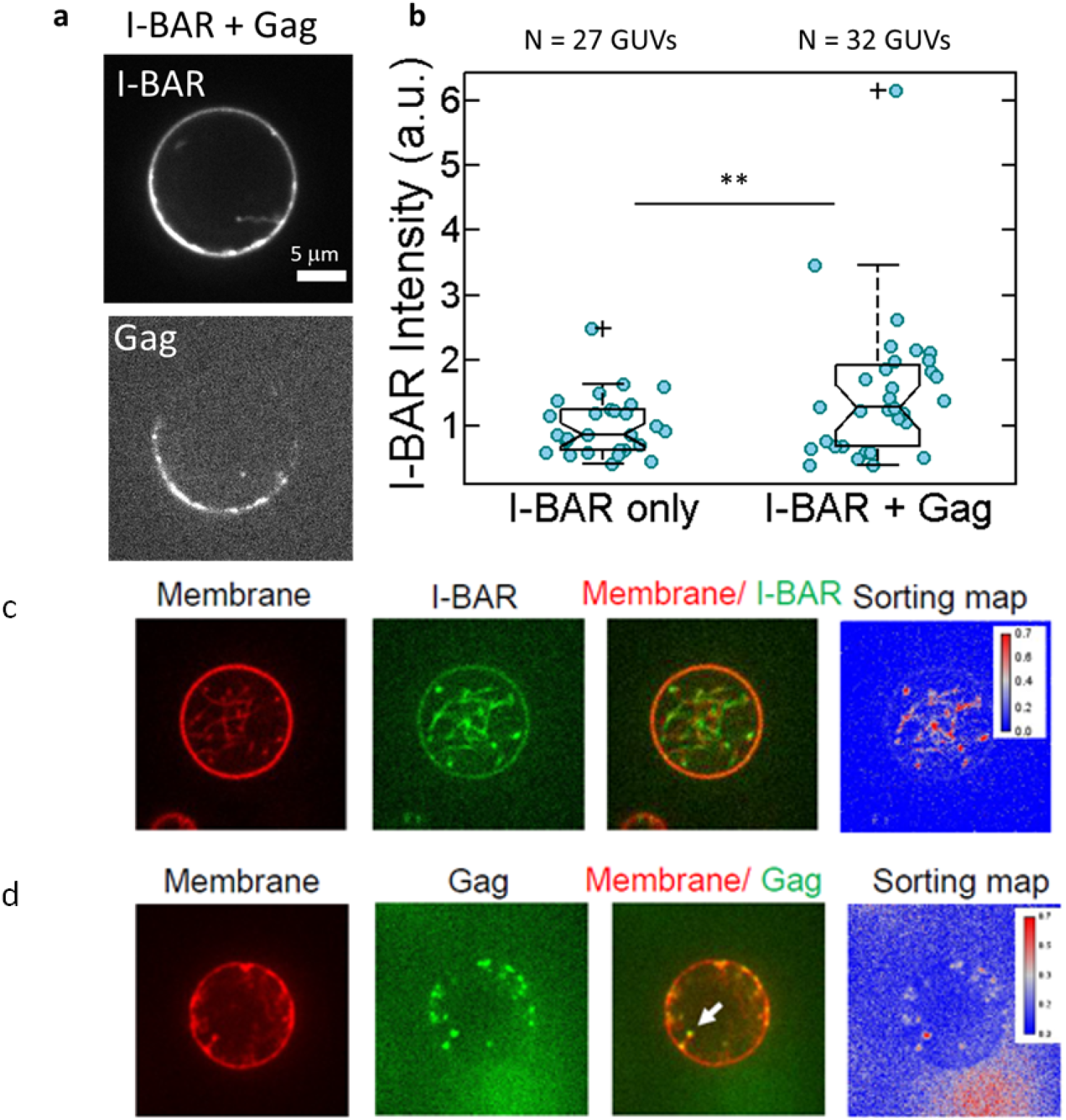
(a,b) In vitro GUV experiments showing an increase of IRSp53 I-BAR binding on GUV upon HIV-1 Gag addition. (a) Representative confocal images of AX488 labelled IRSp53 I-BAR and AX546 labelled Gag on GUV membranes in “I-BAR + Gag” condition. (b) Measurement of AX488 IRSp53 I-BAR fluorescence intensity on membranes in the absence of Gag (named “I-BAR only”), in the presence of Gag where GUVs were first incubated with IRSp53 I-BAR domain and then with HIV-1 Gag (named “I-BAR + Gag”). Each circle presents one GUV analysis. N = 27 GUVs, n = 2 sample preparations for “I-BAR only”, N = 32 GUVs, n = 2 sample preparations for “I-BAR + Gag”. To pool all data points from the 2 sample preparations, I-BAR intensities were normalized by the mean I-BAR intensity in the “I-BAR only” condition. Protein bulk concentrations: 0.05 μM for I-BAR domain and 0.3 μM for Gag. **p = 0.01133, Student’s t-test. **(c,d) In vitro GUV experiments showing incubation of low concentration of the IRSp53-I-BAR domain (green) with GUVS (red)** inducing long membrane tubulation towards the interior of the vesicle (panel-c) while addition of HIV-1 Gag after incubation of I-BAR results in the formation of shorter tubules with HIV-1 Gag sorting to the tip of the short tubules (panel-d).

